# Bazooka/Par3 cooperates with Sanpodo for the assembly of Notch signaling clusters following asymmetric division of *Drosophila* sensory organ precursor cells

**DOI:** 10.1101/2021.01.19.427226

**Authors:** Elise Houssin, Mathieu Pinot, Karen Bellec, Roland Le Borgne

**Affiliations:** Univ Rennes, CNRS, IGDR (Institut de Génétique et Développement de Rennes) - UMR 6290, F-35000 Rennes, France; Equipe Labellisée Ligue Nationale contre le cancer; Wolfson Wohl Cancer Research Centre, Institute of Cancer Sciences, University of Glasgow, Glasgow G61 1QH, UK

## Abstract

In multiple cell lineages, Delta-Notch signaling regulates cell fate decisions owing to unidirectional signaling between daughter cells. In *Drosophila* pupal sensory organ lineage, Notch regulates pIIa/pIIb fate decision at cytokinesis. Notch and Delta that localize apically and basally at the pIIa-pIIb interface, are expressed at low levels and their residence time at the plasma membrane is in the order of the minute. How Delta can effectively interact with Notch to trigger signaling from a large plasma membrane remains poorly understood. Here, we report that the signaling interface possesses a unique apicobasal polarity with Par3/Bazooka localizing in the form of nano-clusters at the apical and basal level. Notch is preferentially targeted to the pIIa-pIIb interface where it co-clusters with Bazooka and the Notch cofactor Sanpodo. Clusters whose assembly relies on Bazooka and Sanpodo activities, are also positive for Neuralized, the E3 ligase required for Delta-activity. We propose that the nano-clusters act as snap buttons at the new pIIa-pIIb interface to allow efficient intra-lineage signaling.

## Introduction

Notch is the receptor of an evolutionarily conserved cell-cell signaling pathway controlling fate acquisition in numerous developmental processes throughout metazoan development (Kopan and Ilagan, 2009). Within many cell lineages, following the division of a precursor cell, Notch activation regulates binary fate choice between daughter cells (Bertet et al., 2014; Bivik et al., 2016; Dong et al., 2012; Ohlstein and Spradling, 2007; Pardo-Saganta et al., 2015; San-Juan and Baonza, 2011). In the majority of cases, Notch receptor is activated by transmembrane ligands present in adjacent cell. Following binding to Notch, endocytosis of the ligand generates pulling forces driving a change in the conformation of the Notch extracellular domain leading to the exposure of the S2 cleavage site of Notch (Gordon et al., 2015; Langridge and Struhl, 2017; Meloty-Kapella et al., 2012; Seo et al., 2016; Shergill et al., 2012; Wang and Ha, 2013). This mechanosensitive cleavage is followed by a constitutive proteolytic cleavage of Notch by the gamma secretase complex (Mumm et al., 2000; Struhl and Adachi, 2000) giving rise to the transcriptionally active Notch intracellular domain (NICD) (Kopan and Ilagan, 2009). Since proteolytic activation of the Notch receptor is irreversible, Notch activation needs to be tightly controlled in time and in space. However, the spatio-temporal cascade of the events remains poorly characterized.

Sensory organ precursors (SOPs) of the pupal notum of *Drosophila* have been instrumental to study intra-lineage, Notch-dependent fate decisions (Schweisguth, 2015). SOPs are epithelial cells that divide asymmetrically within the plane of a single-layer epithelium to generate two daughter cells, an anterior pIIb cell and a posterior pIIa cell, precursors of internal and external cells of the sensory organ, respectively. The pIIa-pIIb fate acquisition relies on the differential activation of Notch during cytokinesis as a result of the unequal partitioning of Numb and Neuralized (Neur) in pIIb cell (Le Borgne and Schweisguth, 2003; Rhyu et al., 1994). In pIIb, Numb interacts with Sanpodo (Spdo), a four pass transmembrane protein interacting with Notch, regulating its trafficking and required for Notch signaling activation (Babaoglan et al., 2009; Cotton et al., 2013; Couturier et al., 2013; Johnson et al., 2016; Upadhyay et al., 2013). Numb causes the targeting of Spdo/Notch to late endosomes (Cotton et al., 2013; Couturier et al., 2013), whereas Neur promotes the endocytosis of the Notch ligand Delta (Dl) (Le Borgne and Schweisguth, 2003), so that Notch is inhibited in pIIb and activated in pIIa. During SOP cytokinesis two pools of Notch, located apical and basal to the midbody, are present at the pIIb-pIIa interface and both contribute to Notch signaling (Bellec et al., 2021; Trylinski et al., 2017). Previous studies based on photobleaching and photoconversion experiments revealed that the basal pool of Notch is the main contributor of NICD (Trylinski et al., 2017). However, how the two pools of Notch are targeted along the pIIa-pIIb interface to promote this private intra-lineage cell-cell communication rather than with the neighboring epidermal cells remains largely unknown.

In vertebrates, the scaffolding protein Par3 has been reported to regulate Numb-mediated trafficking of Integrin and Amyloid precursor protein (APP). Indeed, by binding to the phosphotyrosine domain of Numb, Par3 precludes Numb from binding to Integrin, thus hindering Numb from causing Integrin endocytosis (Nishimura and Kaibuchi, 2007). Similarly, Par3 interferes with the interaction between Numb and APP. In the absence of Par3, there is an increase in Numb-APP interaction leading to decreased surface APP and increased targeting of APP to late endosomal-lysosomal compartments (Sun et al., 2016). Whether Numb and Bazooka (Baz), the *Drosophila* ortholog of Par3, are interfering one with each other to control Spdo/Notch trafficking in SOP daughters is unknown.

In addition to regulating Numb-mediated membrane trafficking Par3 regulates adherens junction (AJs) organization and forms a complex with Par6 and atypical Protein Kinase C (aPKC), a complex that is essential in establishment or maintenance of epithelial cell apicobasal polarity (Assemat et al., 2008; Goldstein and Macara, 2007; Laprise and Tepass, 2011; Nelson, 2003; Rodriguez-Boulan and Macara, 2014; St Johnston and Ahringer, 2010). During SOP mitosis, the unequal segregation of Numb relies on the SOP-specific remodeling of polarity modules and the phosphorylation by the Baz-aPKC-Par6 complex (Bellaiche et al., 2001a; Wirtz-Peitz et al., 2008). Prior to mitotic entry, aPKC-Par6 are in complex with the tumor suppressor Lethal Giant Larvae (Lgl). Assembly of the Baz-aPKC-Par6 complex is initiated upon phosphorylation of Par6 by the mitotic kinase AuroraA (AurA), then causing the autoactivation of aPKC. aPKC next triggers the phosphorylation of Lgl. Phosphorylated aPKC and Par6 can assemble with Baz (Wirtz-Peitz et al., 2008). The Baz complex localises at the posterior apical and lateral cortex, while Discs-large (Dlg) and Partner of Inscuteable (Pins) accumulates at the anterior lateral cortex during SOP mitosis (Bellaiche et al., 2001b; Roegiers et al., 2001). Baz complex phosphorylates

Numb at the posterior cortex, thereby preventing Numb to localise there, resulting in the unequal distribution of Numb in the anterior cortex (Smith et al., 2007; Wirtz-Peitz et al., 2008). Following degradation of AurA at metaphase to anaphase transition, Baz may be released from the Par6-aPKC complex. The localisation and potential functions of Baz versus aPKC/Par6 complex during cytokinesis of SOP, as well as the consequence of the polarity remodeling at mitosis on the apico-basal polarity of the pIIa-pIIb interface at the time of Notch activation are unknown.

In this study, we have analysed the remodeling of cell-cell junction markers and polarity determinants throughout SOP cytokinesis and compared it to that of epidermal cell cytokinesis. We report that in the SOP, the PAR complex is dismantled during cytokinesis with aPKC redistributing in intracellular apical compartments, while Baz localises into apical and lateral clusters along the pIIa-pIIb interface together with Notch and Spdo. Analyses of clone borders revealed that Notch is not uniformly distributed at the plasma membrane but is instead selectively enriched at the pIIa-pIIb interface, indicative of a polarized transport mechanism towards the SOP daughter cells interface. Baz and Spdo, but not Notch, are required for the formation of the clusters. Neur localised in the clusters while Numb prevents cluster occurrence. We propose a model according which Baz regulates Notch and Spdo localisation into clusters to favor signaling.

## Results

### Remodeling of apico-basal polarity during SOP division leads to an atypical pIIa-pIIb interface

As SOP undergoes a specific remodeling of polarity modules during division (Bellaiche et al., 2001a), we started by investigating the resulting apicobasal polarity and remodeling of junctional complexes on the forming pIIa-pIIb cell interface during cytokinesis from which Notch is activated. We previously reported that formation of the novel adhesive pIIa-pIIb interface, visualized with DE-Cadherin-GFP (E-Cad), is assembled with similar kinetics to that of epidermal daughters (Founounou et al., 2013). Here, we first live-monitored and compare the remodeling of septate junctions (SJs) and markers in SOP versus epidermal cells as they undergo cytokinesis. All fluorescent markers are inserted at the locus, giving rise to functional reporters expressed at physiological level. In every case, the transition from metaphase to anaphase was considered to be t0 (time is indicated in min:sec). Dlg-GFP (Woods and Bryant, 1991) and Neuroglian-YFP (Nrg-YFP; (Genova and Fehon, 2003)), two markers of SJs, are progressively recruited at the new pIIa-pIIb interface, immediately basal to the AJs around 25 min after anaphase, with similar kinetics but with a lower intensity than in epidermal daughters (Fig. S1A-A’ Nrg-YFP: t23±6 min at pIIa-pIIb *versus* t24±4 min at epi-epi; Fig. S1B-B’ Dlg-GFP: t27±6 min at pIIa-pIIb *versus* t24±4 min at epi-epi interface, and data not shown). The Nrg-YFP signals at the interface with epidermal cells spread more basally compared to surrounding epidermal cells (Fig. S1A, A’, see orthogonal sections at t6). While the morphological differences possibly reflect differences in the para-cellular diffusion barrier composition and/or organisation, assembly of AJs and SJs occurs with similar kinetics along the new pIIa-pIIb and epidermal cell interfaces.

We next analysed the localisation of the component of the subapical complex Crumbs (Crb,) as well as two members of the Par complex: aPKC and Baz. Crb-GFP is detected faster at the new apical pIIa-pIIb interface than between epidermal daughters (Fig. 1A, S1C; t7±2 min at pIIa-pIIb interface *versus* t13±3 min at epi-epi interface). Then, while Crb remains localised at apical interface of epidermal cells (Fig. S1C), in pIIa and pIIb cells, Crb-GFP localised primarily in apical cytoplasmic puncta (t13±4 min; Fig. 1A and S1C’) at the expense of the pIIa-pIIb cell interface (t23±4 min). A similar behaviour was observed for aPKC-GFP (Besson et al., 2015) that is first localised at the new pIIa-pIIb interface (t7±1 min at pIIa-pIIb interface (Fig. 1B and S1D’), *versus* t12±3 min at epi-epi interface (Fig. S1D)) and then redistributed in part to cytoplasmic puncta primarily in the pIIa cell. In striking contrast to aPKC and Crb, Baz-GFP presents a unique pattern of localisation (Fig. 1C). First, it is not relocalised in apical cytoplasmic puncta. Second, Baz-GFP is localised both at the pIIa-pIIb interface and enriched at the posterior pole of the pIIa cell (Fig. 1C, t9 min) when compared to epidermal-epidermal daughter cell interface (Fig. S1E, t12 min, upper panels) in agreement with previous reports (Le Borgne et al., 2002; Roegiers et al., 2001). Third, Baz-GFP signal is more punctiform along the apical pIIa-pIIb interface than at the interface between two epidermal cells (Fig. 1C and S1E, upper panels). Last, Baz-GFP localises in clusters at the lateral pIIa-pIIb interface (Fig. 1 C, C’ middle panels and orthogonal views, see also Supplemental Movie 1). These clusters that are specific to SOP daughter cells interface appear concomitantly to the first apical punctiform localisation, ~10 min after the onset of anaphase (Fig. 1 C, C’).

**Figure 1:**
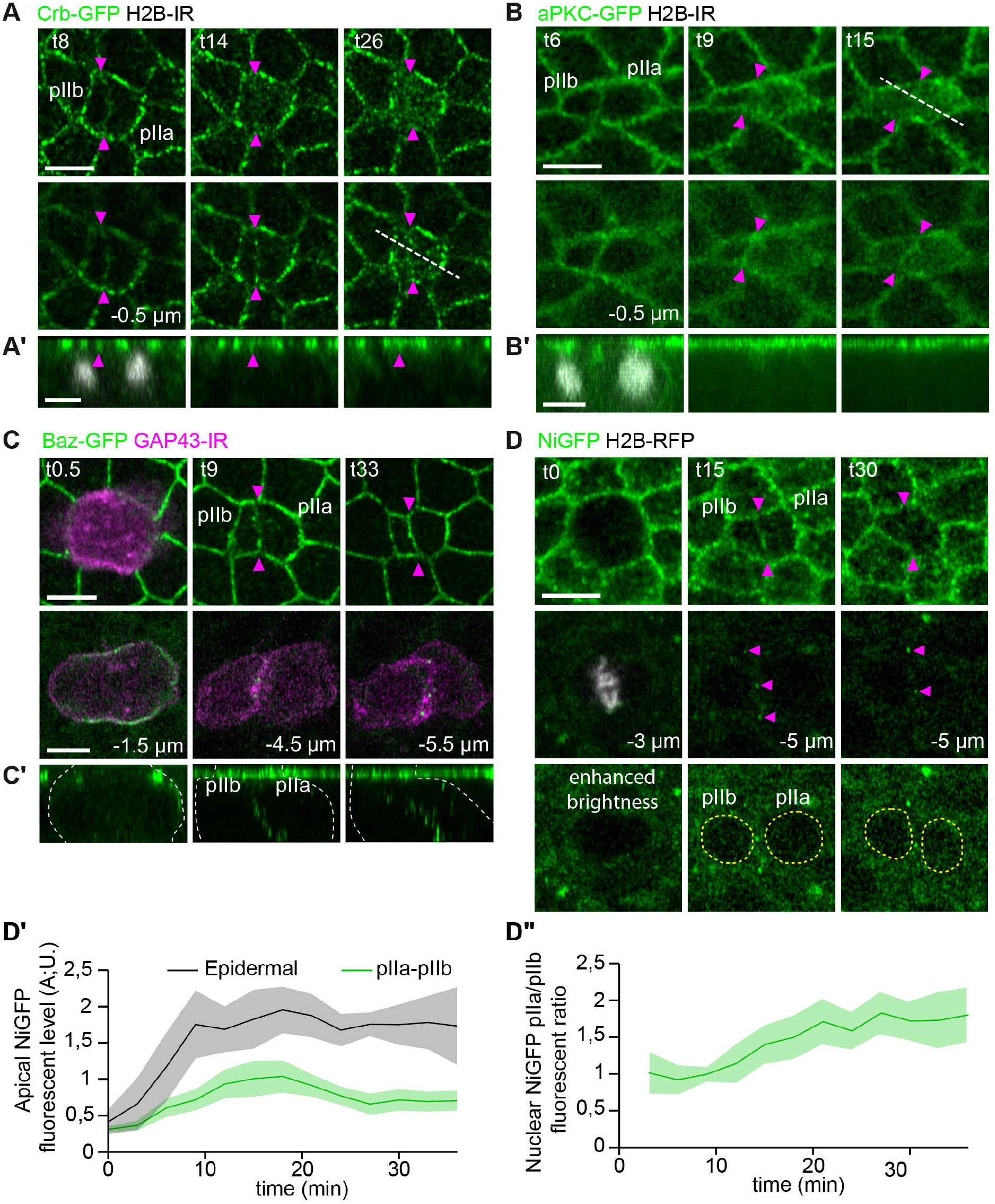
Distribution of polarity markers and Notch during SOP cell division. Time-lapse imaging of Crb-GFP (**A,A’**, n=22 SOPs), aPKC-GFP (**B, B’**, n=10 SOPs), Baz-GFP (**C, C’**, n=28 SOPs) and NiGFP (**D**, n=25 SOPs) during SOP cytokinesis that gives rise to the anterior pIIb and the posterior pIIa daughter cells. (**A-B’**). Crb-GFP and aPKC-GFP are first localised to the pIIa/pIIb interface and then re-localise into punctiform structures in the medial part of the cell at t26 and t15, respectively. The white line at t26 (**A**) and t15 (B) are used for the plot profiles presented in Fig S1 C’, D’, respectively. (**C-C’**) Apically, Baz-GFP signal is more punctiform and enriched at pIIa/pIIb interface as well as on pIIa posterior membrane compared to surrounding epidermal interfaces. Laterally Baz-GFP is found in clusters along the pIIa/pIIb interface (**D**). NiGFP localises transiently at the apical pIIa-pIIb interface (t15, magenta arrowheads, top panels) and in lateral clusters at the level of the nuclei (magenta arrowheads, middle panels). The SOPs and their daughter cells were identified by H2B-IRF670 (**A-B**), GAP43-IR (**C**), or H2B-RFP (**D**) expressed under the *neur* minimal promoter. (**A-C**) and (**A’-C’**) depict top views and corresponding orthogonal views along the new pIIa-pIIb interface delineated by the magenta arrowheads. The t0 corresponds to the onset of anaphase transition. In **C’**, the white dashed line delineates the SO cell boundaries. (**D’**) Quantitation of the NiGFP signal presents at the apical pIIa-pIIb interface (green) or epidermal daughter cell interface (grey) (**D”**) and of the nuclear NiGFP signal (ratio pIIa/pIIb nuclei, delineated by yellow dashed lines in **D**) over time post anaphase onset. Time is in min, scale bars are 5 μm.

We concluded that the apico-basal polarity is specifically remodelled during SOP cytokinesis giving rise to a pIIa/pIIb interface with an atypical polarity, raising the possibility that polarity reshaping may play a role in Notch receptor localisation and activation that we next investigated.

### Polarity remodeling of the SOP is concomitant with localisation of Notch at the pIIa-pIIb interface

The distribution of Baz is reminiscent of that reported for a GFP tagged version of Notch, NiGFP, that transiently distributes in clusters at the apical and lateral pIIa-pIIb interface prior to proteolytic activation and targeting of the resulting intracellular domain of Notch tagged with GFP into the nucleus of the pIIa cell (Fig. 1D,D”, (Bellec et al., 2018; Bellec et al., 2021; Couturier et al., 2012; Trylinski et al., 2017). As Baz, Notch is not detected in clusters at the lateral interface of epidermal daughter cells (Fig. S1F). As epithelial cells are tightly packed, we first determined the origin of the Notch signal presents between the SOP daughter cells. Indeed, during epithelial cells cytokinesis, the dividing cell maintains membrane contacts with the neighboring cells forming a *ménage à quatre* that is progressively resolved as the cell progresses towards abscission (Daniel et al., 2018; Founounou et al., 2013; Guillot and Lecuit, 2013; Herszterg et al., 2013; Morais-de-Sa and Sunkel, 2013; Wang et al., 2018). Because of its duration, the contact is particularly noticeable within the plane of SJs where epidermal cells maintain contact in the form of fingers like protrusions connected to the SOP midbody (t5, Fig. S1B) until the entire belt of SJ is reformed (about 90 min) (Daniel et al., 2018). Analysis of the borders of clones of cells expressing NiGFP (Bellec et al., 2018) enables to determine the origin of the NiGFP signal detected along the pIIa-pIIb interface (Fig. 2A-B’). When a SOP expressing NiGFP is dividing next to epidermal cells expressing untagged Notch (Fig. 2A), NiGFP signal is detected at the apical pIIa-pIIb interface, and basally between the pIIa-pIIb nuclei (Fig. 2A’, t21). In the converse situation, when a SOP expressing untagged Notch is dividing next to epidermal cells expressing NiGFP, no GFP signal is detected at the apical and basal pIIa-pIIb interface (Fig. 2B,B’). Analyses of clone boundaries also revealed that low NiGFP signal was detected at the boundary between SOP daughters and the neighboring epidermal cells. This is not observed in epidermal cell where Notch equally partitions along the plasma membrane (Fig. 2A’, B’, and Fig 3A). These data show that, following SOP division, Notch is preferentially transported towards or stabilized at the pIIa-pIIb interface where signaling takes place.

**Figure 2:**
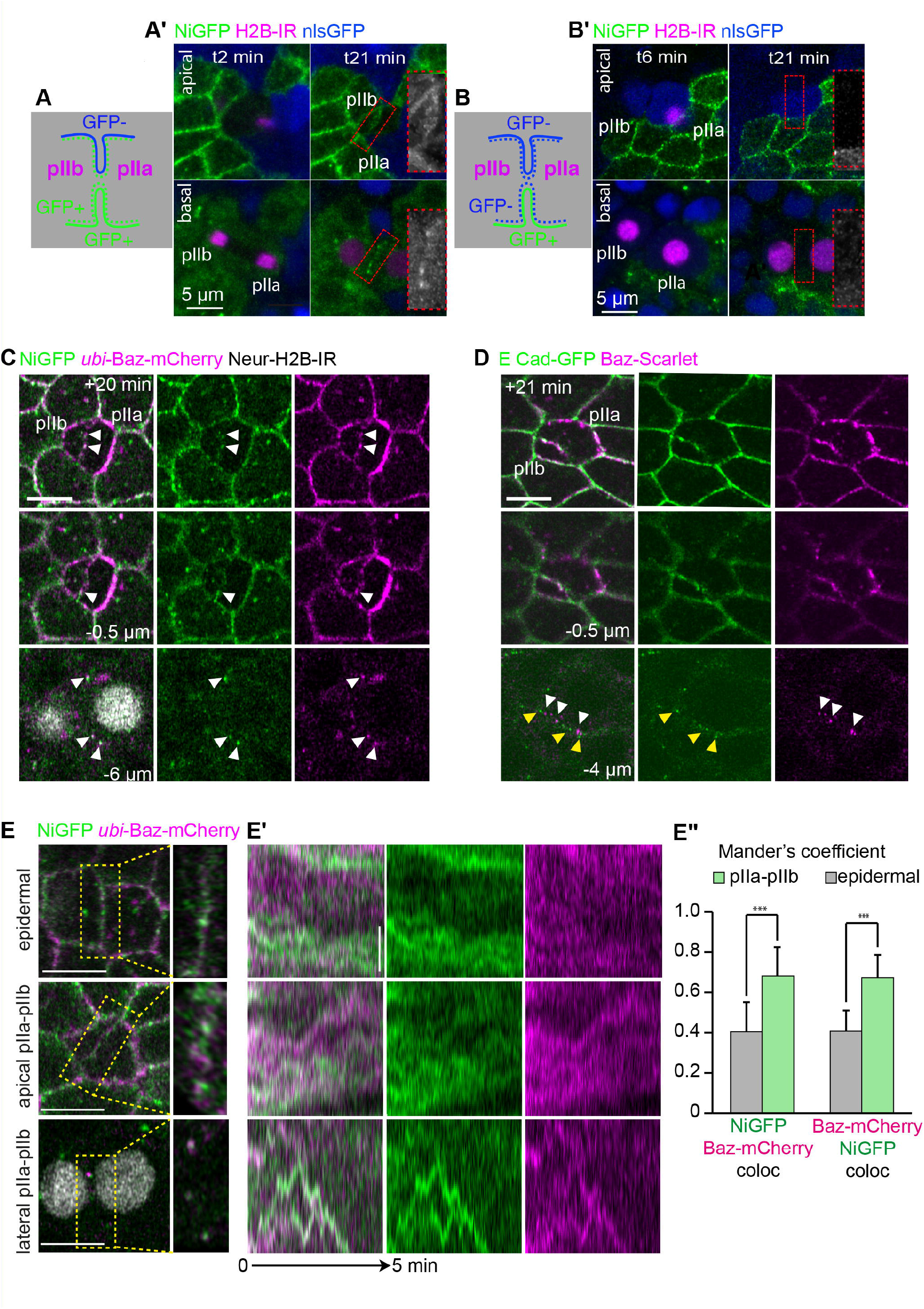
Dynamics of colocalisation of NiGFP and Baz-mCherry at pIIa-pIIb interface. **(A-B’)** Schematic representations **(A, B)** and original data **(A’, B’)** illustrating borders of clones of cells expressing NiGFP adjacent of cells expressing untagged versions of Notch. The dividing SOP expresses NiGFP in (**A, A’**) while SOP NiGFP-negative is adjacent to NiGFP-positive cells in (**B, B’**). Dashed lines and continuous lines in (**A**) and (**B**) represent the plasma membrane of the SOP daughters and that of epidermal cells, respectively. At t21, NiGFP signal localizes at the pIIa-pIIb apical interface and in lateral clusters (delineated by a red dashed rectangle in **A’** versus **B’**) showing that this signal comes from the SOP daughters, not from the neighboring epidermal cells. **(C)** Live imaging of NiGFP together with *ubi*-Baz-mCherry in SOP daughter cells (n=9 SOPs). Baz colocalises with NiGFP in apical and lateral clusters (white arrowheads). **(D)** Live imaging of E-Cad-GFP (green) together with Baz-Scarlet (magenta) during SOP cytokinesis. Baz colocalises in part with E-Cad at the apical pIIa-pIIb interface, whereas the Baz-positive lateral clusters (white arrowheads) are largely distinct form the E-Cad lateral clusters (yellow arrowheads). In addition, in contrast to Baz-positive lateral clusters, E-Cad lateral clusters are not restricted to the pIIa-pIIb interface (n=9 SOPs). SOPs and their daughter cells were identified by H2B-IR **(A’, B’, C, D and E)** expressed under the *neur* minimal promoter. **(E,E’)** Kymographs generated from high temporal resolution acquisitions (every 2 seconds) of epidermal-epidermal interface or pIIa-pIIb interface, taken from the insets highlighted with yellow dashed lines in panel (**E**) (n=10 for each type of cell?). Apically, acquisitions were taken around +12 min after anaphase and laterally around +20 min after anaphase. On the kymographs, tracks correspond to the movement of the clusters. We can see a colocalisation of NiGFP and Ubi-Baz-mCherry only at pIIa/pIIb interface apically as well as laterally. (**E”**) Mander’s colocalisation coefficient measured on the kymographs realised from apical or lateral acquisitions (**E**). Manders’ coefficients are significantly higher for pIIa/pIIb interface kymographs compared to epidermal interface kymographs. ***p-value ≤ 0.001. Time is in min, scale bars are 5μm in **A’, B’, C** and **D** and 1μm in **E.**

As Notch resembles Baz localisation at the pIIa-pIIb interface, we investigated their localisation simultaneously by co-imaging NiGFP together with Ubi-Baz-mCherry (Bosveld et al., 2012). We first observed that Ubi-Baz-mCherry colocalises with NiGFP in punctae along the pIIa-pIIb apical interface as well as in the basally located, lateral clusters (Fig. 2C). The NiGFP/Baz lateral clusters do not correspond to spot AJs (McGill et al., 2009) as E-Cad and Baz do not colocalise at the lateral pIIa-pIIb interface (Fig. 2D). We next monitored the dynamics of Ubi-Baz-mCherry and NiGFP clusters using high spatiotemporal resolution acquisitions. Kymographs of these acquisitions (Fig. 2 E,E’) show that, at the apical pIIa-pIIb interface and even more markedly at the lateral interface, the Ubi-Baz-mCherry and NiGFP tracks colocalise to a greater extent compared to epidermal-epidermal interface (Fig. 2E-E”), raising the possibility that Baz and Notch act together in space and time to contribute to pIIa/pIIb identities that we next investigated.

### Baz contributes to Notch localisation after SOP division

After having establish that Ubi-Baz-mCherry/NiGFP clusters are specific to the pIIa-pIIb interface, we asked whether Notch and Baz are mutually required for clusters formation. In *baz^EH747^* clones, a genetic and protein null allele of Baz ((Shahab et al., 2015), Fig. S2A,A’), NiGFP is still detectable in clusters along the pIIa-pIIb interface (Fig. 3A, A’). However, the apical NiGFP fluorescence intensity measured as described in (Bellec et al., 2021) is significantly reduced (Fig. 3B) and the numbers and intensity of the lateral clusters are decreased upon loss of Baz (Fig. 3C, C’). Similar decrease in the number and intensity of Notch clusters was observed upon Baz silencing (Fig. S2B-C’). Despite the fact that the nuclear pIIa-pIIb ratio of NiGFP, used as a proxy for Notch activation, is on average similar for *baz^EH747^* mutants and controls (Fig. 3B’), the decrease of cluster numbers and fluorescence intensity is accompanied by defective Notch activation in 3 to 9% of the *baz^EH747^* homozygous mutant SO (Fig. S2D). Collectively, these results indicate that Baz contributes to Notch localisation and, albeit not strictly required, modulates activation of Notch signaling in the pIIa cell.

**Figure 3:**
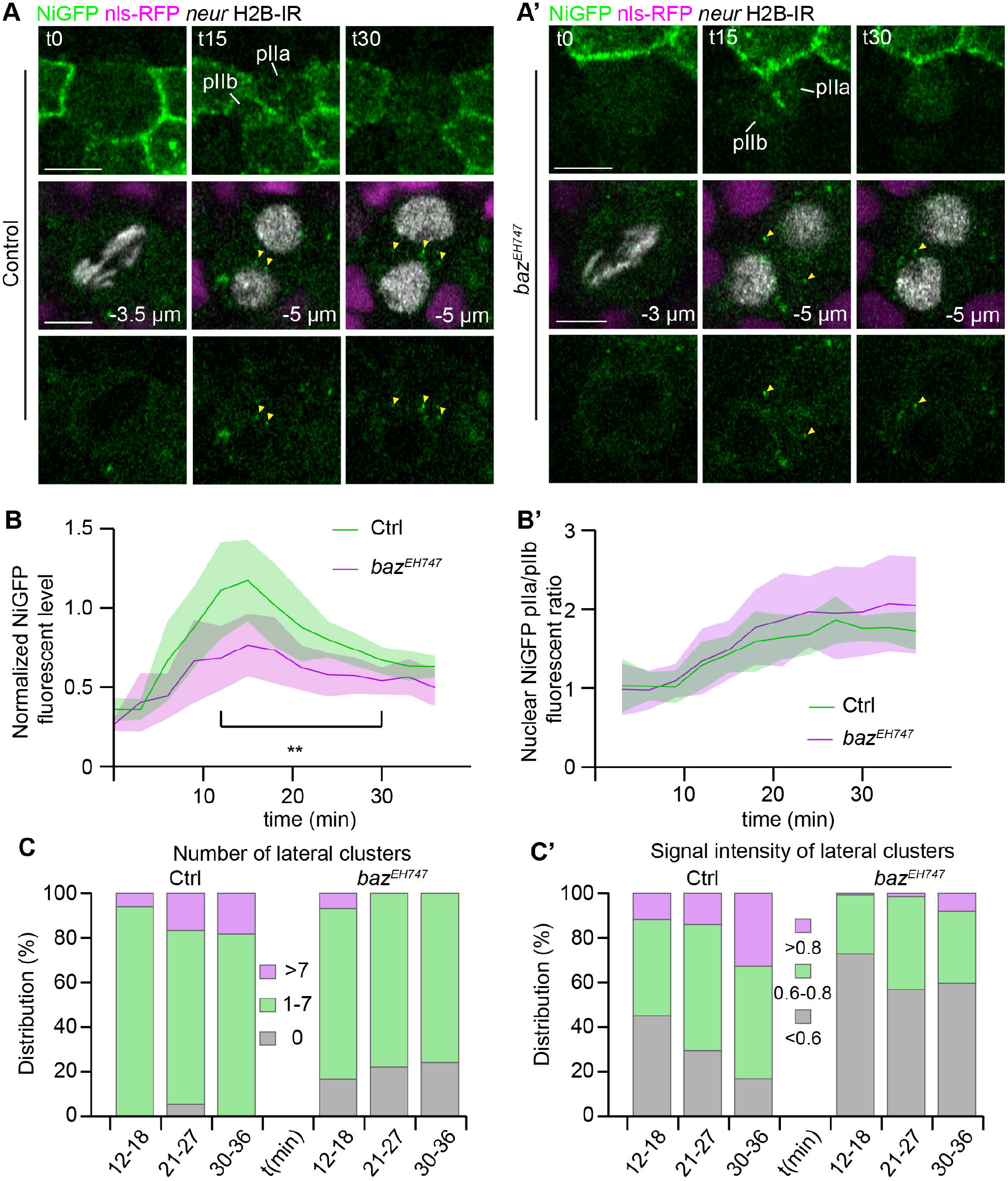
NiGFP localisation in *baz^EH747^* mutant clones. (**A, A’**) Live imaging of NiGFP in control (**A**) or *baz^EH747^* mutant clones (**A’**) clones (ctrl, n=9; *baz^EH747^*, n=10). The localisation of NiGFP in *baz^EH747^* mutant clones is similar to the control with a transitory recruitment to the apical pIIa/pIIb interface and the presence of lateral clusters (yellow arrowheads). (**B**) Quantification of NiGFP fluorescence intensity at the apical pIIa-pIIb interface (ctrl, n=9; *baz^EH747^*, n=10). The apical fluorescent level measured is significantly lower in *baz^EH747^* mutant clones compared to ctrl. (**B’**) Nuclear NiGFP fluorescent level ratio between pIIa and pIIb over time after SOP anaphase (ctrl, n=9; *baz^EH747^*, n=10). The ratio increases progressively in both *baz^771747^* mutant clones and ctrl reaching similar values. Time is in min, scale bars are 5 μm. **(C,C’)** Histogram representing the evolution of the number of clusters **(C)** and fluorescence intensity of the clusters (**C’**) over time in ctrl (n=8) and *baz^EH747^* (n=10) mutant clones. Whereas clusters number and intensity tend to slightly increase over time in ctrl, in *baz^EH747^* mutant clones, clusters number and intensity are quite stable over time and tend to be lower compared to ctrl.

To test if conversely Notch is required for Baz localisation in clusters, we depleted Notch using RNAi or degradFP system (Caussinus et al., 2011). Both approaches resulted in the reduction in Notch signal and Notch *loss-of-function* phenotypes, including an excess of SOP specification due to defective lateral inhibition, and pIIa to pIIb cell fate transformation ((Fig. 4 A-C) and data not shown). Under these conditions, Baz still localises in clusters along the apico-basal pIIa-pIIb interface as in the wild type, indicating that Notch is dispensable for Baz cluster assembly (Fig. 4 D, E, yellow arrowheads). Nonetheless, lateral Baz clusters were never detected between two adjacent SOPs nor between a SOP prior division and a pIIb-pIIb-like cell (data not shown). These data suggest the remodeling of cell polarity taking place during SOP cytokinesis is a prerequisite for the assembly of Baz/Notch lateral clusters.

**Figure 4:**
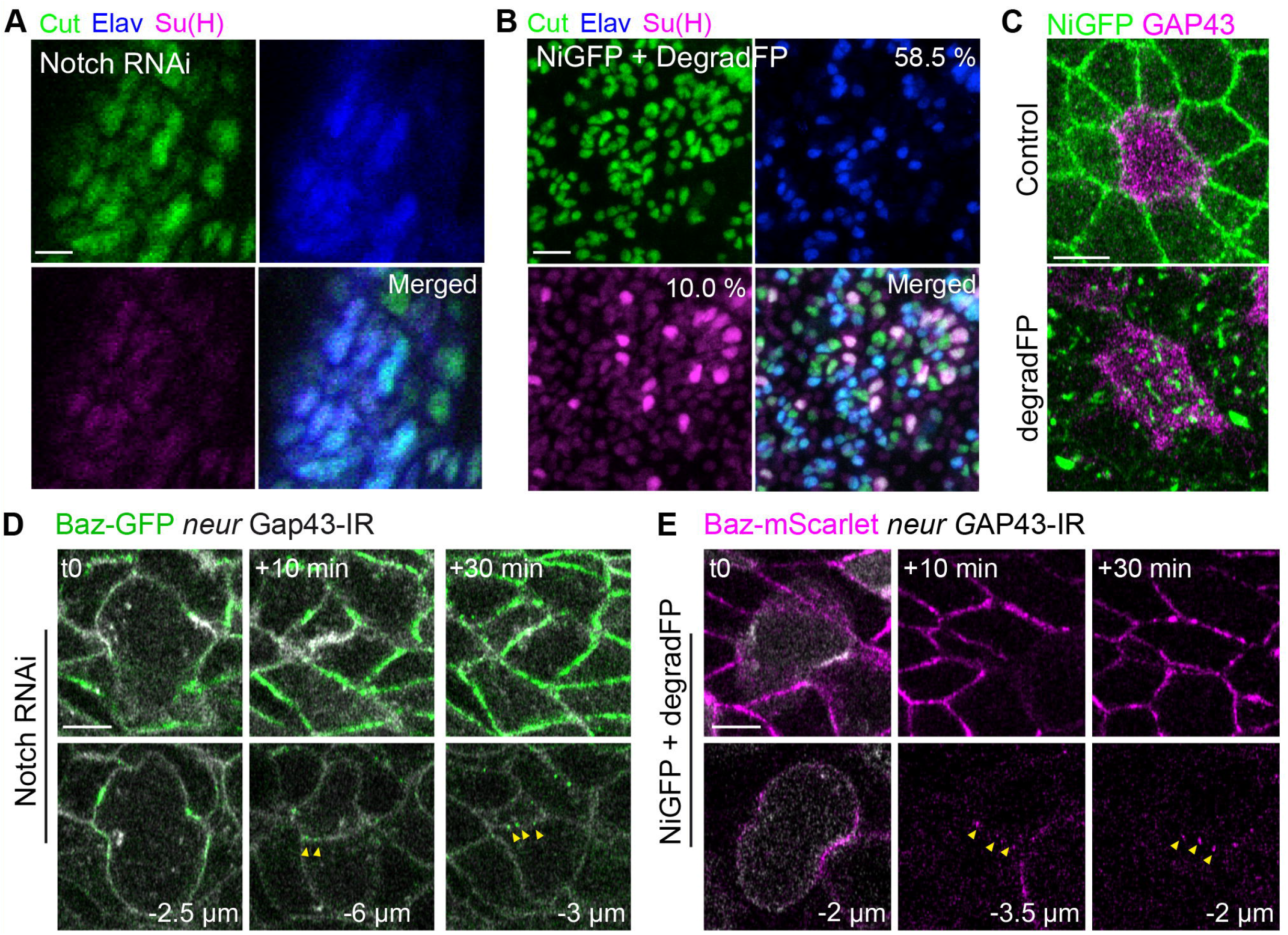
Notch *loss-of-function* does not impair the formation of Baz clusters along the pIIb-pIIb-like interface. (**A-B**) SO lineage analyses using the SO marker Cut (green), the socket marker Su(H) (magenta) and the neuronal marker Elav (blue) upon silencing of Notch (**A**) and depletion of NiGFP with degradFP (**B**). In Notch RNAi, neurons were largely present among all the Cut+ cells, at the expense of Su(H) cells (**A**, n=2 nota). In NiGFP + degradFP (**B**), there are 58.5% of neurons for only 10% of socket-positive cells (n=1045 Cut+ cells, n=4 nota). (**C**) Localization of NiGFP (green) together with *neur*-GAP-43-IR (magenta) in control and with degradFP. Upon degradFP based depletion of NiGFP nota, NiGFP cortical signal is drastically reduced compared to control, and mainly bright intracellular GFP-positive punctae are observed which could correspond to partially degraded NiGFP (n=19). (**D-E**) Localisation of Baz-GFP (green in **D**) and Baz-Scarlet (magenta in **E**) together with *neur*-GAP43-IR upon Notch depletion using RNAi (**D**) or degradFP (**E**). Following Notch depletion, Baz still localises apically and in lateral clusters at the pIIa-pIIb interface (yellow arrowheads). Time is in min. Scale bars are 5 μm.

As Baz was deemed necessary but not sufficient *per se* for Notch cluster assembly, we next asked what key regulators of Notch-dependent binary fate acquisition control Baz/Notch cluster assembly and/or dynamics.

### The assembly of Baz-Notch clusters at pIIa-pIIb interface is modulated by Numb, Neur and Delta

To further investigate the requirements for the presence of Baz-Notch clusters at the pIIa-pIIb interface, we first investigate the localisations and functions of regulators of Notch signaling during asymmetric cell division, starting with the cell fate determinant Neur. As previously reported, Neur-GFP (Perez-Mockus et al., 2017b) localises asymmetrically at the anterior cortex, opposite to Baz during SOP prometaphase and is unequally partitioned in pIIb cell (Fig. 5A). At t21min, Neur is localised in clusters at the pIIa-pIIb interface where it largely colocalised with Baz-Scarlet (Fig. 5A and insets). Loss of Neur results in a Notch *loss-of-function* phenotype and an increased transient signal of NiGFP at the apical pIIa-pIIb interface (Fig. 5 B, C) accompanied with a higher number and brighter lateral clusters of NiGFP (Fig. 5B, C’, C”), that persisted more than 36 min after anaphase onset. Similar results were obtained upon silencing of Delta (Fig S3 A-C’). Upon loss of Neur, Baz localises at the apical interface and in clusters at the lateral pIIa-pIIb interface (Fig. 5D). By analyzing their dynamics at high spatio-temporal resolution, we found that Baz and NiGFP remain closely associated in the apical and lateral clusters at the pIIa-pIIb interface upon loss of Neur (Fig. 5E). Collectively, these results argue that Neur, Delta, Baz and Notch localise together in clusters at the pIIa-pIIb interface and that their numbers and signal intensities depend on Neur and Delta activities. These data also suggest that the clusters are assembled but fail to dissociate in the absence of Neur-mediated activation of Notch.

**Figure 5:**
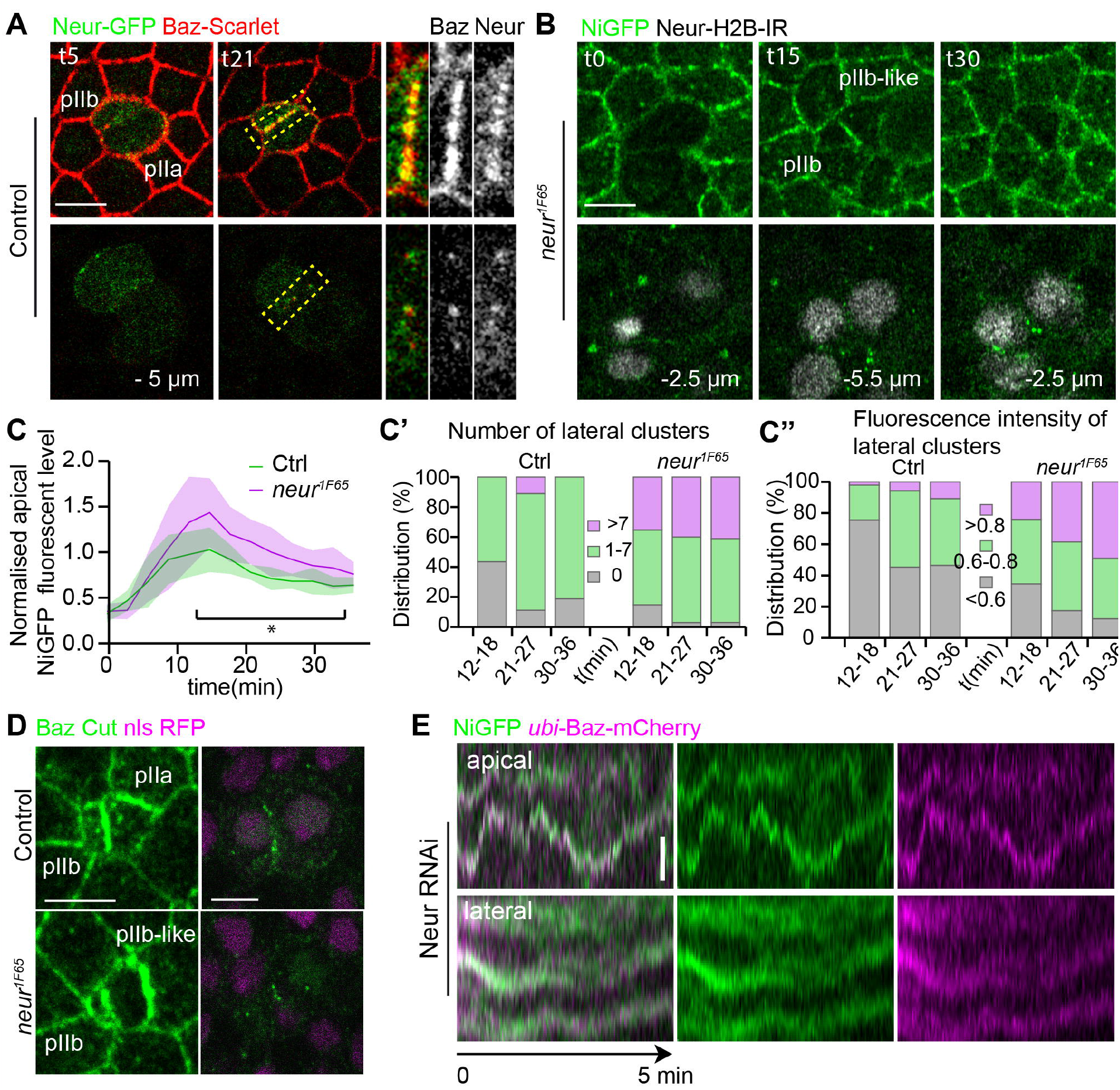
Neur localises in and regulates the number of NiGFP/Baz positive clusters. (**A**) Localization of Neur-GFP (green) and Baz-Scarlet (red) in control. Neur and Baz colocalise in clusters both apically and laterally at the pIIa-pIIb interface (n= 8 SOPs). Yellow dashed line delineates the enlarged areas presented in the insets (right panels). **(B-C”)** Time lapse imaging of NiGFP (green) together with *neur* H2B-IR (grey) in *neur^1F65^* (**B**) (n=9 for control, and n=10 for *neur^1F65^*). Loss of Neur results in increased amounts of NiGFP at the apical pIIa-pIIb interface (t15 in **B**, quantitation in **C**)), and an increase in the number and signal intensity of lateral clusters (t30 in **B**), a phenotype quantitated in (**C’,C”**). (**D**) Localisation of Baz (green) in control and in *neur^1F65^*. Baz localises in apical and lateral clusters upon loss of Neur as in the control. In **B** and **D**, *neur^1F65^* mutant cells were identified by the loss of nlsRFP (n=9, 7 nota for control; n=13, 7 nota for *neur^1F65^* condition). (**E**) kymographs illustrating the colocalisation of NiGFP (green) together with Ubi-Baz-mCherry (magenta) over time upon depletion of Neur (n=11 SOPs). Time is in min, scale bars are 5 μm and 1 μm for the kymographs.

We next analysed the localisation and function of Numb in the occurrence of Notch-Baz clusters. In contrast to Neur (Fig. 5A), Numb does not colocalise with Baz-Scarlet-positive clusters at the pIIa-pIIb interface (Fig. 6A). Silencing of Numb results in a Notch *gain-of-function* phenotype with increased transient signals of Notch at the apical pIIa-pIIa like interface (Fig. 6B-C), accompanied with more numerous and brighter lateral clusters of NiGFP (Fig. 6B, C’, C”). These data are fully consistent with that published previously in (Couturier et al., 2012; Trylinski et al., 2017) and further show that the lateral Notch clusters accumulation persists until at least 36 min after anaphase onset. Upon silencing of Numb, Baz localises in lateral clusters at the pIIa-pIIa like interface (Fig. 6D, D’) where it colocalises with NiGFP as revealed by the dynamics of the Baz-Notch clusters at high spatio-temporal resolution (Fig. 6E).

**Figure 6:**
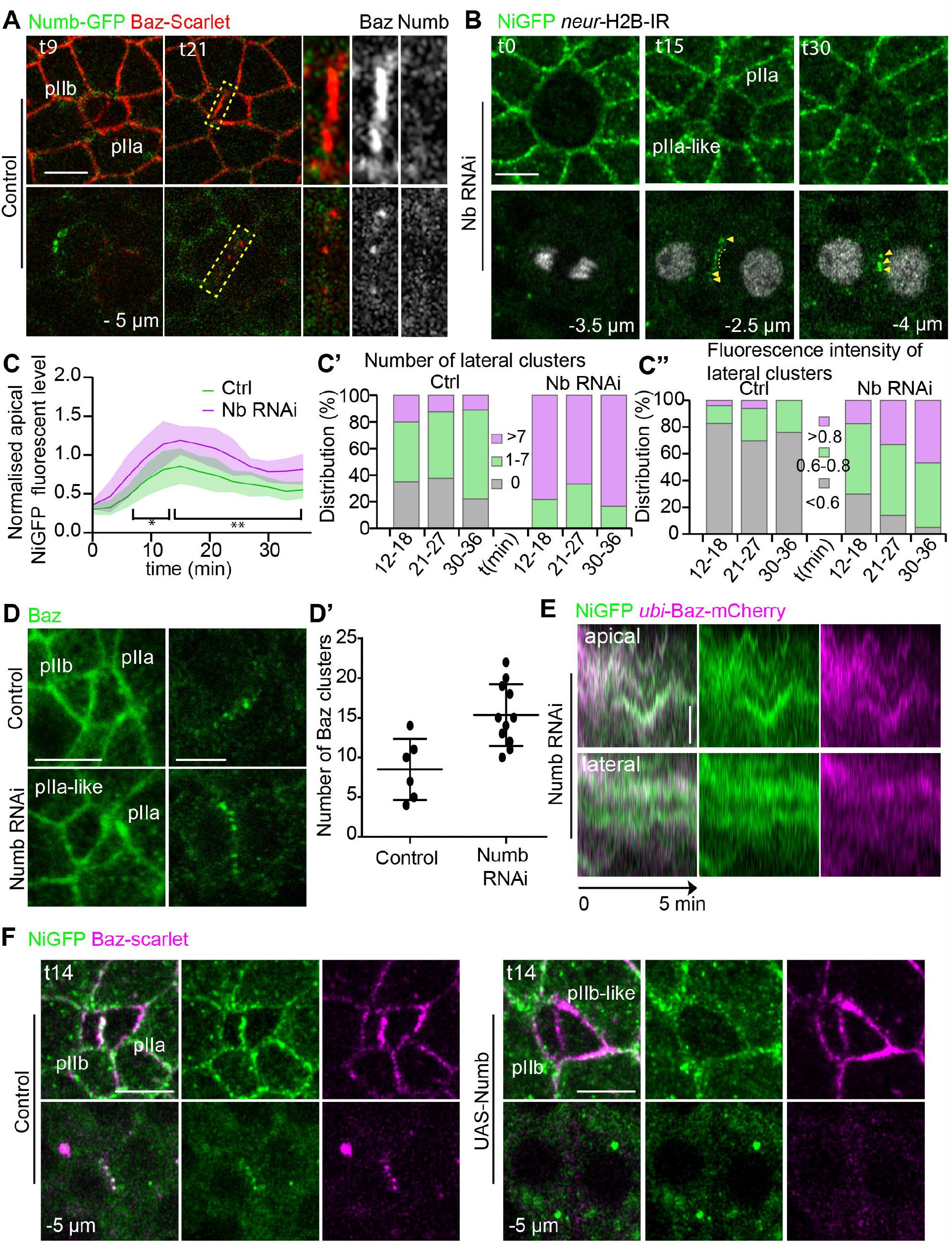
Numb is absent from but regulates the number of NiGFP/Baz positive clusters. (**A**) Localisation of Numb-GFP (green) and Baz-Scarlet (red) in control. Numb and Baz do not colocalise in clusters apically nor laterally (insets corresponding to dashed yellow rectangles) at the pIIa-pIIb interface (n= 5). **(B-C”)** Time lapse imaging of NiGFP (green) together with *neur* H2B-IR (grey) upon silencing of Numb (**B**) (n= 20 for NumbRNAi). Loss of Numb results in increased amounts of NiGFP at the apical pIIa-pIIb interface (t15 in **B**, quantitation in **C**), and an increase in the number and signal intensity of lateral clusters (t30 in **B**), a phenotype quantitated in **C’** and **C”**, respectively. Yellow arrowheads and dashed yellow line point to lateral clusters. (**D, D’**) Localisation of Baz (green) in control and in Numb RNAi. Baz localises in the apical and lateral clusters upon depletion of Numb, as in the control (n=15 SOPs, 5 nota). (**D’**) Quantitation of the number of Baz clusters along the pIIa-pIIb interface. (**E**) Kymographs illustrating the colocalisation of NiGFP (green) together with Ubi-Baz-mCherry (magenta) over time upon silencing of Numb (n= 14 SOPs). **(F)** Time lapse imaging of NiGFP (green) together with Baz-Scarlet (magenta) in control (left panel, n=14) and upon overexpression of Numb (right panels, n=8 SOPs). Overexpression of Numb results in the loss of NiGFP signal at the pIIa-pIIb interface both apically and laterally compared to control. Time is in min, scale bars are 5 μm and 1 μm for the kymographs.

Together, these data indicate that Numb represses Notch activation and negatively regulates the number of Notch-Baz clusters. As Numb is present and regulates Notch endosomal trafficking in the anterior pIIb cell (Cotton et al., 2013; Couturier et al., 2013), our data suggest that Notch-Baz clusters are assembled in the anterior cell Plutot a l’interface non? Pas de data montrant que c’est dans la cellule anterieure upon loss of Numb Mais aussi en absence de Neur ou Delta, and contribute to Notch activation in this cell. This model further suggests that Numb acts antagonistically to Baz to promote Notch clusters assembly and/or stability Neur et Delta aussi non?. To test this prediction, we overexpressed Numb in the SOP and daughter cells and observed that NiGFP is no longer detected along the pIIb-pIIb-like interface neither apically nor laterally (Fig. 6F). While Baz localises uniformly at the apical SOP daughter cell interface, lateral clusters are barely detectable (t14, Fig. 6F, bottom panels). Thus, Numb appears to inhibit Baz-Notch cluster laterally, raising the possibility that Numb and Baz could act antagonistically as proposed in vertebrates (Nishimura and Kaibuchi, 2007; Sun et al., 2016). Du coup, je crois avoir deja pose la question, mais si on surexprime Neur ou Delta, a t on le meme phenotype? Quelle est le modele que vous avez en tete les concernant? As Numb interacts with the NPAF motif of Spdo to control Notch/Spdo endosomal trafficking, the above data are calling the question of the relationship between Baz and Spdo that we next studied.

### Spdo is required for Baz-Notch clusters formation

On live specimens Baz-Scarlet and Spdo-GFP (Couturier et al., 2013) colocalized both at the apical pIIa-pIIb interface as well as in the lateral clusters (Fig. 7A, t21). Compared to the control situation, the Baz-positive lateral clusters are no longer detectable upon loss of Spdo (Fig. 7B and S4C). In agreement with published results (Couturier et al., 2012), loss of Spdo also results in an increase of NiGFP signal at the apical SOP daughters interface and the appearance of a continuous and nebulous staining of NiGFP instead of the characteristic, well defined, lateral clusters observed at the pIIa-pIIb interface of controls SO (Fig. 7C,D, and S4A, A’). We also noticed that NiGFP persists at the apical interface compared to the control, and that NiGFP is detected apically, in the cytoplasm or at the apical plasma membrane, indicative of higher levels of Notch upon loss of Spdo (Fig. 7C, t15 and t30, upper panels). Fluorescence Recovery After Photobleaching (FRAP) analyses reveal that the NiGFP signal at the apical interface is recovered 1.9 time faster, with a mobile fraction 1.6 time higher than in the control situation (Fig. 7 E,E’ and S4B). The changes in NiGFP distribution and time residence at the pIIb-pIIb like interface are accompanied by a loss of colocalisation of NiGFP and Baz-mCherry at the apical and lateral pIIb-pIIb-like interface upon Spdo silencing (Fig. S4 C-D’). We first concluded that Spdo coclusters with Baz and Notch at the pIIa-pIIb interface and second, that the activity of Spdo is required for the clustering of Baz/Notch along the pIIa-pIIb interface to promote Notch activation.

**Figure 7:**
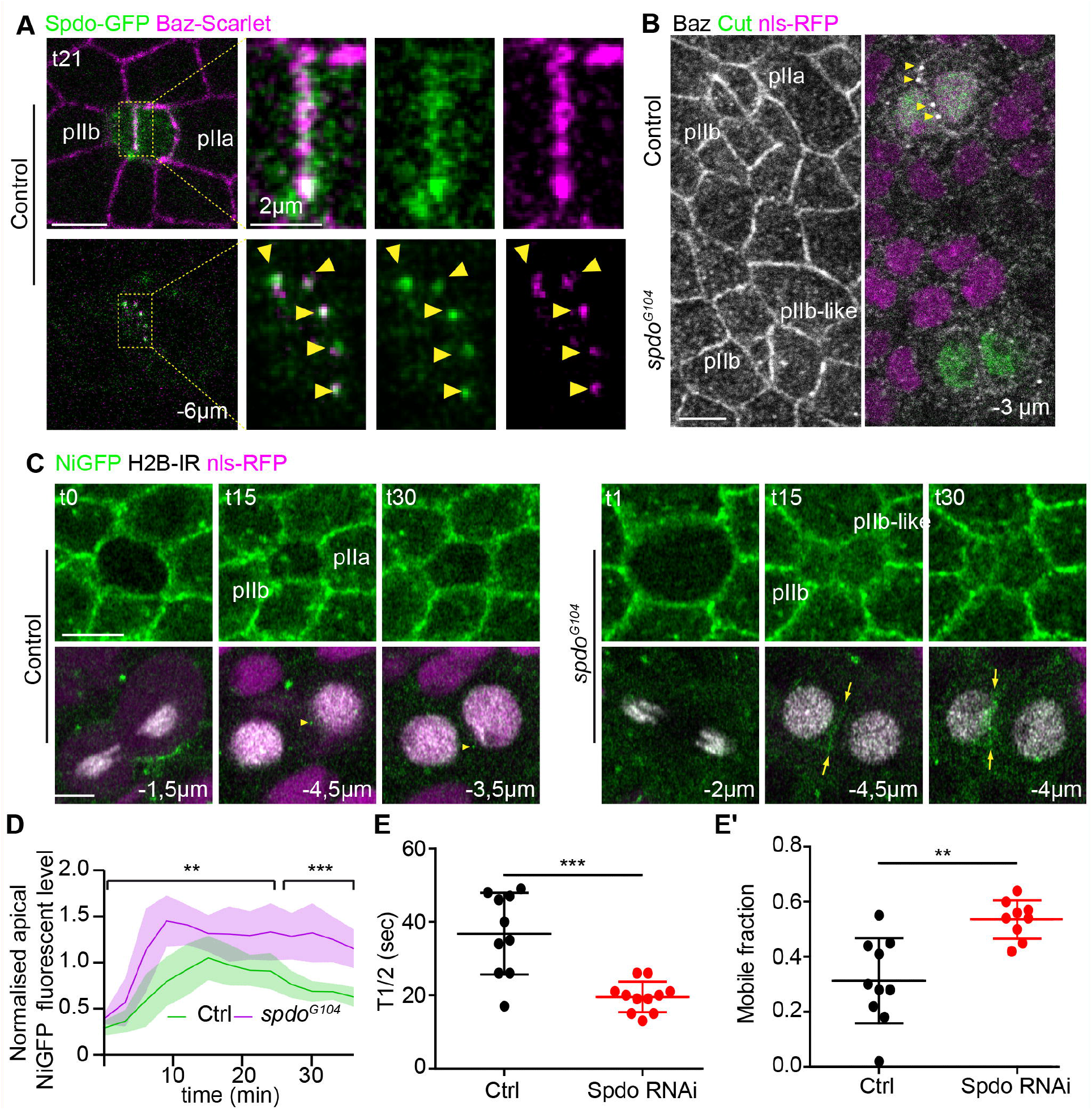
Sanpodo is required for the assembly of NiGFP/Baz. (**A**) Localisation of Spdo-GFP (green) and Baz-Scarlet (magenta) in control. Spdo colocalises with Baz-Scarlet (magenta) in clusters both apically and laterally (yellow arrowheads) at the pIIa-pIIb interface (n= 10, see insets corresponding to dashed yellow rectangles). (**B**) Localisation of Baz (grey) in *spdo^G104^* mutant clones. While Baz localises in clusters at the lateral pIIa-pIIb interface (yellow arrowheads) of control, in *spdo* mutant SO clusters are not detected. Control SOs and *spdo^G104^* mutant cells were identified using Cut (green) and nls-RFP (magenta), respectively. (**C, D**) Time lapse imaging of NiGFP (green) together with *neur* H2B-IR (grey) in control (left panels, n=14) and *spdo^G104^* mutant SO (right panels, n=10 SOPs). SO and *spdo^G104^* mutant SOs were identified using neur H2B-IR (grey) and nls-RFP (magenta), respectively. Loss of Spdo resulted in an increase of NiGFP signal at the pIIa-pIIb interface (quantitation in (**D**). In contrast to the control situation where NiGFP localises in clusters (yellow arrowheads), in *spdo^G104^* mutant SO, NiGFP signal is continuous along the pIIa-pIIb interface (yellow arrows). (**E, E’**) Quantitation of the t1/2 (**E**) and mobile fraction (**E’**) of NiGFP following FRAP at the pIIa-pIIb interface. FRAP analyses reveal that the time residence of NiGFP at the apical interface is 1.9 time shorter upon silencing of Spdo (n=11 SOPs) and the mobile fraction of NiGFP is 1.6 time higher than in control (n= 10 SOPs). Time is in min, scale bars are 5 μm.

## Discussion

In this study, we have characterized the remodeling of apico-basal cell polarity occurring during SOP division leading to a specific pIIa-pIIb Notch signaling interface. We report that Baz, but not aPKC, co-partitions with Notch, Spdo and Neur in apical and lateral clusters. The assembly of these clusters requires Baz and Spdo activities, and their number and dynamics are regulated by Delta, Neur and Numb activities. In the absence of Numb, the number of clusters increases while overexpression of Numb results in their disappearance suggesting that Numb and Baz acts antagonistically. We propose that Notch/Baz/Spdo/Neur clusters represent the Notch signaling units at the pIIa-pIIb interface.

### The pIIa-pIIb interface possesses an atypical apico-basal polarity compared to epidermal cells

Previous pioneer work unraveled that Par3/Par6/aPKC and Pins/Dlg polarity modules are specifically relocated from apical-basal into posterior-anterior cortex respectively during SOP mitosis (Bellaiche et al., 2001a; Roegiers et al., 2001). Assembly of the Baz/Par6/aPKC is initiated by the phosphorylation of Par6 by the mitotic kinase AurA (Wirtz-Peitz et al., 2008). Here, we report that during cytokinesis coinciding with the presumptive proteolytic degradation of AurA, the Baz /Par6/aPKC complex disassembles with aPKC redistributing like Crumbs in apical intracellular compartments at the expense of its regular plasma membrane localisation observed in epidermal cells. In contrast to aPKC, Baz redistributes apically at the posterior pole of the pIIa cell and in the form of clusters at the pIIa-pIIb interface. Such lateral clusters of Baz are only found at the pIIa-pIIb interface indicating that SOP specific remodeling polarity that takes place at SOP mitosis is instrumental in clusters formation. Baz has been reported to be excluded from the lateral plasma membrane following to Par1-mediated phosphorylation (Benton and St Johnston, 2003). In addition, phosphorylation of Baz by Par1 activity is antagonized by type 2A protein phosphatase (PP2A) activity (Krahn et al., 2009), and silencing of *tws* the regulatory B subunit of PP2A results in Notch *gain-of-function* phenotype (Shiomi et al., 1994). However, we observed that Baz lateral clusters are decorated with anti-phospho Baz S151, S980, or S1085 suggesting that while Par1-dependent phosphorylation is taking place efficiently, it does not promote exclusion of Baz from the lateral plasma membrane (data not shown). It is yet unclear how SOP polarity remodeling leads to Baz clusters assembly and localisation laterally. The fact that Spdo and Notch, two transmembrane proteins, colocalise with Baz in the lateral clusters (both on fixed and live specimens) argues against a model according which N-terminal oligomerisation domain of Baz could drive phase separation of Baz (Liu et al., 2020) at this location.

The nanoscopic clusters of Baz are reminiscent to the clusters serving as AJ assembly landmark, by repositioning Cadherin-Catenin clusters at apico-lateral sites for assembly of spot AJ (SAJ, (McGill et al., 2009)). The Baz and Cad-Catenin clusters were shown to assemble independently and the number and size of Cadherin-Catenin clusters are decreased in *baz* mutant as reported here for Notch clusters. This is pointing towards a common function of Baz in the control of cluster assembly, positioning and stability.

In addition to organize membrane nanoscopic clusters of Cadherin and Catenin, in vertebrates, Par3 also functions as a receptor for exocyst, a protein complex of the secretory pathway required for the delivery of basolateral proteins to the plasma membrane (Ahmed and Macara, 2017). It is interesting to note that Baz clusters are exclusively located at the pIIa-pIIb interface. Analyses of NiGFP clones borders revealed a preferential localisation of Notch at the pIIa-pIIb interface instead of being equally partitioned at the plasma membrane. Together with the fact that Sec15, a component of the exocyst complex, regulates Notch and Spdo trafficking to regulate binary fate acquisition in the SO lineage (Jafar-Nejad et al., 2005), our results place Baz as a potential regulator for delivery of Notch/Spdo at specific sites along the pIIa-pIIb interface.

In any case, while Baz activity is required for efficient Notch clusters assembly, only a few cell fate transformations are observed upon loss of Baz. This does not mean that Baz is dispensable for Notch signaling. We propose that Baz activity is required to define a threshold for Notch activation and that Baz *loss-of-function* strongly sensitizes the ability of SOP daughters to signal. Indeed, while the simple silencing of Delta or Baz remains with limited effect on identity acquisition (tufting phenotype and a few pIIa to pIIb transformations for Delta or Baz RNAi respectively), the simultaneous silencing of Baz and Delta leads to a completely bald fly, i.e. a complete loss of Notch function (our unpublished data). Thus, Baz by regulating the size and number of clusters at the pIIa-pIIb interface appears important for proper Notch signaling during SOP cytokinesis.

### Do the Notch/Baz/Spdo clusters constitute signaling units?

The clusters present at the pIIa-pIIb interface are positive for Notch, Spdo, Baz and Neur. While Delta is also detected along the pIIa-pIIb interface on fixed specimen (Bellec et al., 2021), DlGFP was reported to be barely detectable in living pupae unless Neur-mediated Delta endocytosis was blocked (Trylinski et al., 2017). This led to the proposal that newly synthesized Delta reaches the plasma membrane and signals from there thus exhibiting a rapid turnover/endocytosis. A direct implication of these findings is that the clusters are present on both sides of the pIIa-pIIb interface as a kind of snap button with Delta/Neur in the pIIb cell interacting *in trans* with Notch/Spdo in the pIIa cell. Based on the role of Numb in Notch/Spdo trafficking in the pIIb cell, the fact that Baz is enriched in the posterior pIIa cell at cytokinesis and the proposed antagonism between Numb and Baz, we anticipate that Baz is located primarily in the clusters on the pIIa cell side. As the time residence of Delta, Notch and Baz in the cluster are very short (minute time scale), it implies that Delta can interact with Notch *in trans*, be internalized in a Neur-dependent manner to promote the S2 cleavage of Notch in the minute time scale. We propose that Baz mediated clustering might be a mean to locally concentrate Notch/Spdo and increase its ability to interact with Delta.

### Sensory organ specific relocation of Crumbs and site of NICD production

The SO-specific localisation of Crb could be explained by the down-regulation of Expanded, a negative regulator of Crb endocytosis (Besson et al., 2015) and/or by Neur mediated Stardust/Crb endocytosis (Perez-Mockus et al., 2017b; Perez-Mockus and Schweisguth, 2017). In agreement with this latter, expression of Bearded Brd^KR^ (Perez-Mockus et al., 2017a), a negative regulator of Neur results in accumulation of Crb-GFP at the plasma membrane (data not shown). Could the lack of Crb at the pIIa-pIIb interface be a prerequisite for Notch activation? Indeed, in zebrafish, by binding to the extracellular domain of Notch, Crb was reported to inhibit Notch activity (Ohata et al., 2011). In *Drosophila*, Crb is shown to stabilize Notch and Delta at the apical plasma membrane and loss of Crb results in increased Notch signaling (Richardson and Pichaud, 2010). In addition, Crb is reported to negatively regulate gamma-secretase activity, an activity required for the S3 cleavage of Notch (Herranz et al., 2006). Whether the absence of Crb from the apical pIIa-pIIb interface could relieve such inhibitions awaits experimental demonstration. If this would turn out to be the case, this would only apply to apical signaling. Alternatively, as proposed in (Perez-Mockus and Schweisguth, 2017), the Neur-mediated endocytosis of Crb may result in changes in the subcellular distribution of Delta to favor Delta signaling at the basolateral membrane. In any case, our study brings further supports to the notion of a tight coupling between cell polarity and Notch signaling. Photobleaching and phototracking experiments during SOP cytokinesis revealed that among the two pools of Notch, the basolateral pool located basal to the midbody is the main contributor (Trylinski et al., 2017). While the apical pool of Notch also contributes to NICD production, it is yet unclear whether NICD is directly produced from the apical pIIa-pIIb interface or if basolateral relocation is a prerequisite (Bellec et al., 2021; Couturier et al., 2012; Trylinski et al., 2017).

According to that model, NICD production would primarily occur at the lateral pIIa-pIIb interface. Our results showing that the composition of the presumptive signaling clusters are similar at the apical and basolateral pIIa-pIIb interface may indicate that NICD could be directly produced from both sites. The remodeling of cell polarity taking place during SOP cytokinesis could thus enable the formation of equally potent signaling clusters along the pIIa-pIIb interface, favoring private pIIa-pIIb cell-cell communication. The amounts and half-life of such signaling clusters could account for the respective contributions of basal versus apical pools in producing NICD.

### Numb and Baz act oppositely on Notch/Spdo clusters assembly

While loss of Neur and loss of Numb both result to an increase in the number and intensity of Baz/Notch/Spdo clusters, the causes are different. Upon lack of Neur, we anticipate that Delta is bound to Notch in *trans*. In absence of Neur-mediated endocytosis of Delta that exerts pulling force on Notch, the clusters are stabilized/not consumed. Numb interacts physically with Spdo to control the subcellular localisation of the Notch/Spdo complex. In the control situation, Numb is not detected in the Notch/Spdo clusters at the pIIa-pIIb interface, suggesting that Notch and Spdo clusters at the interface are predominantly on the pIIa side. Loss of Numb that leads to recycling of Notch/Spdo towards the plasma membrane of the pIIb cell results in the increase in the number and intensity of Notch/Spdo/Baz clusters at the pIIa-pIIb interface. Overexpression of Numb causes the disappearance of Notch/Spdo clusters at the pIIa-pIIb interface.

By analogy to vertebrates, we anticipate that Numb by its ability to bind to Baz (Nishimura and Kaibuchi, 2007), is somehow competing with Baz for the access to Notch/Sdpo, and therefore formation of Notch/Spdo/Baz signaling clusters. Based on the fact that loss of Spdo leads to a stronger reduction in Baz/Notch clusters assembly, a simple prediction is that Baz interacts with Spdo/Notch. Future studies will aim to investigate this possibility to determine which domain of Baz, its ability to oligomerize in order to promote Notch/Baz/cluster assembly.

### Concluding remarks

Due to the conservation of intra-lineage communication, it will be interesting to investigate whether cell-cell communication interface exhibit an atypical apico-basal polarity and if Par3-dependent clustering of Notch also applies to pass a threshold enabling private communication between daughters in vertebrates.

## Acknowledgements

We thank J. Januschke, F. Schweisguth, A. Wodarz, the Bloomington Drosophila Stock Center, the Vienna Drosophila Resource Center, InDroso and the Developmental Studies Hybridoma Bank for providing fly stocks and antibodies. We also thank the Microscopy Rennes Imaging Center-BIOSIT facility. We thank Emeline Daniel for the initial observations of Baz localisation, Thomas Esmangart de Bournonville and Antoine Guichet for critical reading of the manuscript. This work was suported in part by the ARC (post doctoral fellowship), La Ligue contre le Cancer-Equipe Labellisée (R.L.B.) and the Association Nationale de la Recherche et de la Technologie programme PRC Vie, santé et bien-être CytoSIGN (ANR-16-CE13-004-01 to R.L.B.).

## Author contributions

Conceptualization, E.H. and R.L.B.; Methodology, E.H., M.P., K.B. and R.L.B.; Investigation, E.H., M.P. and R.L.B.; Formal Analysis, E. H. and M.P.; Visualization, E.H., M.P. and R.L.B.; Writing – Original Draft, E.H. and R.L.B.; Writing – Review & Editing, E.H. and R.L.B.; Funding Acquisition, E.H. and R.L.B.; Supervision, R.L.B.

## Material and methods

### Star Methods

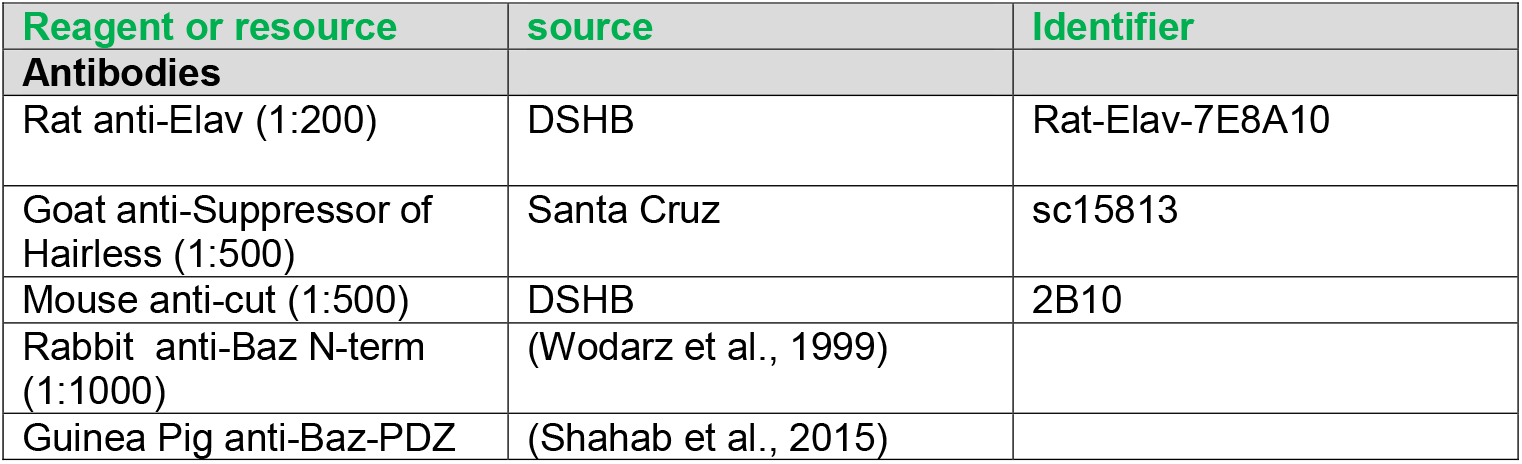
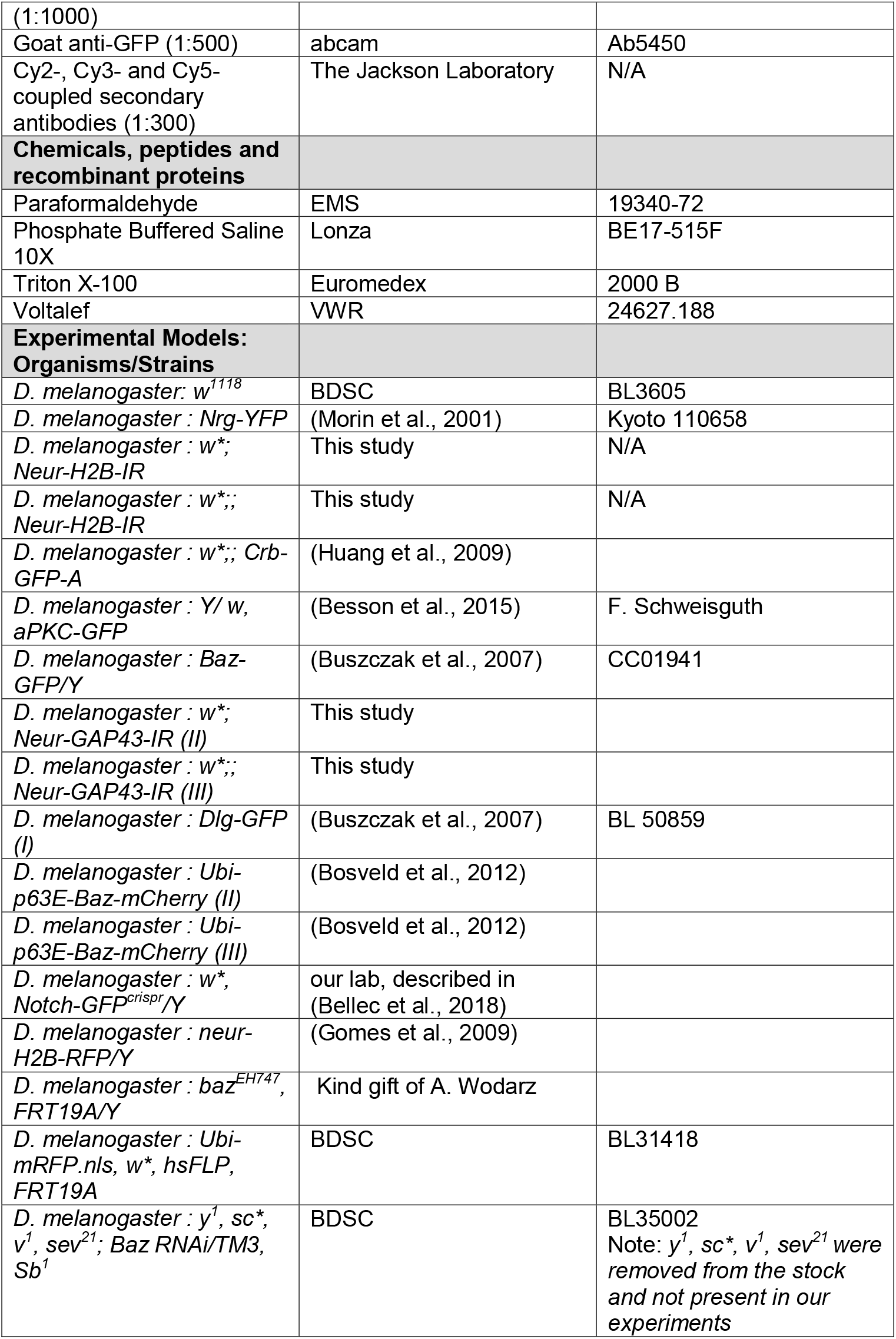
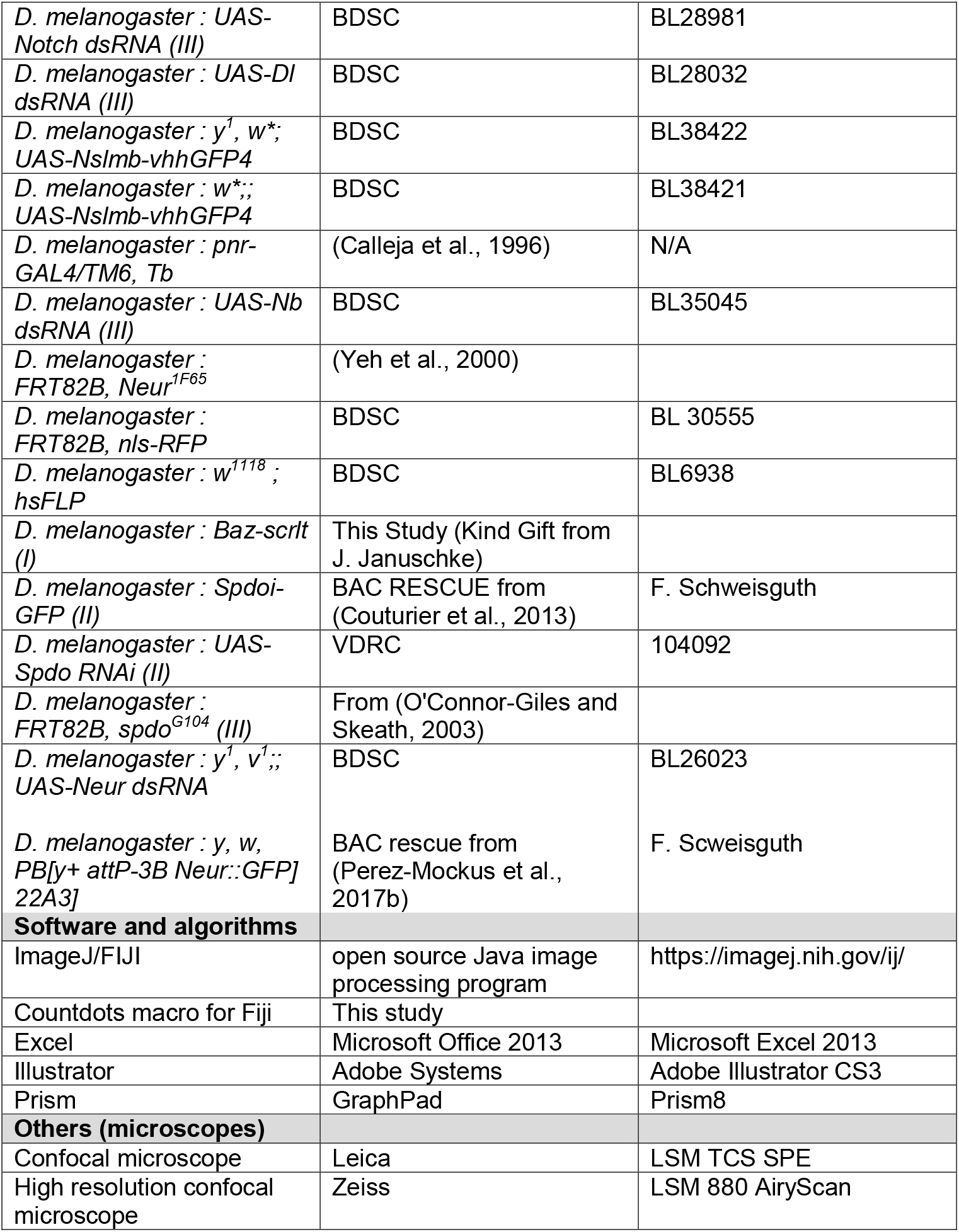

### CONTACT FOR REAGENT AND RESOURCE SHARING

For all kind of requests regarding the material and methods used in this study, please refer to the lead Contact, Roland Le Borgne.

### EXPERIMENTAL MODEL AND SUBJECT DETAILS

#### Drosophila stocks, genetics and CRISPR-mediated homologous recombination (HR)

*Drosophila melanogaster* strains were grown and crossed at 25°C Somatic clones were generated using the FLP-FRT system with a hs-FLP. Crosses were passed in new tubes every 2 days and then, FLP expression was induced by at least 2 heat shocks (1h at 37°C) from embryonic stage for *baz^EH747^* clones and from first instar larval stage for all the other clones.

The *pnr*-GAL4 driver was used to drive the expression of: *UAS-Notch dsRNA; UAS-Nslmb-vhhGFP4 (II and III); UAS-Nb dsRNA; UAS-Spdo dsRNA; UAS-Neur dsRNA and UAS-Dl dsRNA* CRISPR-mediated HR was used to tag the endogenous Baz gene with His-Tag-mScarlet by inDroso (Rennes, France). The Baz-mScarlet resulted from insertion of His-Tag-mScarlet followed by a STLE linker at the amino acid position 40 of Baz (isoforms RA and RC). The gRNA was selected using the Optimal Target Finder tool (http://flycrispr.molbio.wisc.edu/tools), 5’ AAAGCCAAACGCAGGTGAAAAGG, cutting in the second intron (position X:17178549). Complete strategy available upon request to the lead contact).

#### Drosophila genotypes

**Figure 1**

A. *neur*-H2B-IR; Crb-GFP/Crb-GFP
B. aPKC-GFP/Y; *neur*-H2B-IR/+
C. Baz-GFP/Y;; *neur*-GAP-43-IR/+
D. NiGFP/Y; *neur*-H2B-IR/+

**Figure 2**

A. NiGFP, FRT19A/*ubi*-mRFP-nls, w*, *hs*-FLP, FRT19A; *neur*-H2B-IR/+
B. NiGFP, FRT19A/*ubi*-mRFP-nls, w*, *hs*-FLP, FRT19A; *neur*-H2B-IR/+
C. NiGFP/Y; *ubi*-Baz-mCherry/*neur*-H2B-IR;
D. Baz-mScarlet/Y; E Cad-GFP/+
E. NiGFP/Y; *ubi*-Baz-mCherry/Neur-H2B-IR

**Figure 3**

NiGFP, FRT19A/*ubi*-mRFP.nls, w*, hsFLP, FRT19A;; Neur-H2B-IR/+
A’. NiGFP, *baz^EH747^*, FRT19A/*ubi*-mRFP.nls, w*, hsFLP, FRT19A;; Neur-H2B-IR/+

**Figure 4**

A. and D. Baz-GFP/Y;; UAS-Notch RNAi/*pnr*-GAL4
B, C. NiGFP, Baz-mScarlet/Y; UAS-Nslmb-vhhGFP4/+; *pnr*-Gal4/*neur*-GAP43-IR
E Baz-mScarlet/Y; UAS-Nslmb-vhhGFP4/+; *pnr*-Gal4/*neur*-GAP43-IR

**Figure 5**

A. Baz-mScarlet/Y;; attP(Bac Neur-GFP) 22A3 (Perez-Mockus et al., 2017b) /+
B. NiGFP/Y; *neur*-H2B-IR/*hs*-FLP; FRT82B, nls-RFP/FRT82B, *neur^1F65^* C-C”, Ctrl: NiGFP, *neur*-H2B-RFP;; *pnr*-GAL4/+ NiGFP/Y; *neur*-H2B-IR/*hs*-FLP; FRT82B, nls RFP/FRT82B, *neur^1F65^ NiGFP/Y or X; neur-H2B-IR/hs-FLP; FRT82B, nls RFP/FRT82B, neur^1F65^*
D. *hs*-FLP; FRT82B, nls-RFP/FRT82B, *neur^1F65^*
E. NiGFP/Y; *ubi*-Baz-mCherry/Neur-H2B-IR; *pnr*-GAL4/Neur RNAi

**Figure 6**

A. Baz-mScarlet/Y; Numb-GFP (Bellec et al., 2018)/ +
B-C”, Ctrl: NiGFP, *neur*-H2B-RFP;; *pnr*-GAL4/+ NiGFP, *neur*-H2B-RFP;; *pnr*-GAL4/UAS-Nb RNAi
D. *pnr*-GAL4/UAS-Nb RNAi
E. NiGFP/Y; *ubi*-Baz-mCherry/*neur*-H2B-IR; *pnr*-GAL4/Numb RNAi
F. control NiGFP, Baz-mScarlet/Y;; *pnr*-GAL4/+ NiGFP, Baz-mScarlet/Y;; *pnr*-GAL4/Numb RNAi

**Figure 7**

A. Baz-mScarlet/Y; +:+; SpdoiGFP (Couturier et al., 2013)/+
B. NiGFP, /Y; *neur*-H2B-IR/hs-FLP; FRT82B, nlsRFP/FRT82B, *spdo^G104^*
C. NiGFP/Y; *neur*-H2B-IR/hs-FLP; FRT82B, nlsRFP/FRT82B, s*pdo^G104^*
D. E, control: NiGFP/Y; *neur*-H2B-IR; *pnr*-GAL4/+ NiGFP/Y; *neur*-H2B-IR; *pnr*-GAL4/Spdo RNAi

#### Supplemental Figures

**Figure S1**

A. Nrg-YFP/Y; *neur*-H2B-IR/+
B. Dlg-GFP/Y; *ubi*-Baz-mCherry/neur-H2B-IR; *neur*-GAP43-IR/+
C. *neur*-H2B-IR; Crb-GFP/Crb-GFP
D. aPKC-GFP (Besson et al., 2015)/Y; *neur*-H2B-IR/+
E. Baz-GFP/Y;; *neur*-GAP-43-IR/+
F. NiGFP/Y; *neur*-H2B-IR/+

**Figure S2**

A,A’ and D. *baz^EH747^*, FRT19A/*ubi*-mRFP.nls, w*, hsFLP, FRT19A;; Neur-H2B-IR/+
B-C’. control: NiGFP/Y, *neur*-H2B-RFP;; *pnr*-GAL4/+ NiGFP/Y, *neur*-H2B-RFP;; *pnr*-GAL4/RNAi Baz

**Figure S3**

A-C’ control: NiGFP:Y, *neur*-H2B-RFP; +:+; *pnr*-GAL4/+ NiGFP/Y, *neur*-H2B-RFP; RNAi Delta/+; *pnr*-GAL4/+

**Figure S4**

A,A’, and B. NiGFP/Y; *neur*-H2B-IR; *neur*-GAP43-IR, *pnr*-GAL4/Spdo RNAi
C. NiGFP, Baz-mScarlet/Y;; *pnr*-GAL4/Spdo RNAi
D,D’. control NiGFP/Y; *ubi*-Baz-mCherry/Neur-H2B-IR; *pnr*-GAL4/+ NiGFP/Y; *ubi*-Baz-mCherry/Neur-H2B-IR; *pnr*-GAL4/Spdo RNAi

## METHOD DETAILS

### Immunofluorescence

Pupae aged of around 17h after puparium formation (APF) were dissected in Phosphate-Buffered Saline (PBS, pH 7.4) and fixed 15 min in 4% paraformaldehyde at room temperature. They were then permeabilized performing 3 washes of 3 min in PBS + 0.1% Triton X-100 (PBT) and incubated with the primary antibodies (in PBT) for 2 h at room temperature or O/N at 4°C. After 3 washes of 5 min in PBT, pupae were incubated for 1h with the secondary antibodies (in PBT). Samples were then washed 3 times in PBT and 1 time in PBS and finally mounted in 0,5% N-propylgallate, 90% glycerol in PBS 1X. After at least 45 min in the mounting medium, images were acquired on a LSM TCS SPE and processed using FIJI.

### Live-imaging and image analyses

Pupae aged of around 16h30 APF were prepared for imaging as described previously (Daniel et al., 2018). Briefly, the pupa is positioned between a glass slide and a coverslip coated with a thin layer of Voltalef, the coverslip being supported anteriorly and posteriorly by columns made of 5 and 6 little coverslips. Images were acquired at 25°C on a LSM 880 AiryScan or LSM TCS SPE and processed using FIJI.

### QUANTIFICATION AND STATISTICAL ANALYSIS

#### Statistical tests

Statistical difference between two conditions was evaluated by a F test followed by a student t test using Microsoft Excel. Statistical significances were represented as follows: *: p value ≤ 0.05; **: p value ≤ 0.01; ***: p value ≤ 0.001.

#### Fluorescent level measurement and analysis

NiGFP apical fluorescence level at the new pIIa/pIIb interface was measured using Fiji (version 1.52) on a z-projection summing 3 slices separated by 0.5 μm. A line of 30 pixels width was traced across the pIIa/pIIb interface to generate a kymograph on which another line of 20 pixels width was drawn all along the time. A *Plot Profile* then gave us the fluorescent levels (in a.u.) for each time point. These values were then corrected for the bleaching over time. To do this, on the same z-projection we measured the fluorescent level of 3 different areas around the SOP, calculated the apical mean fluorescence and determined a bleaching correction factor (t0 apical mean fluorescence/ti apical mean fluorescence) for each time point that we applied to the previous measurements at the new pIIa/pIIb interface. Finally, we normalised to the t0 apical mean fluorescence. For the pIIa/pIIb nuclear fluorescent ratio, we first determined the 3 most centred slices for each pIIa and pIIb nucleus at each time point. We then measured the fluorescent levels on the z-projection summing the determined slices in round areas with constant size and shape positioned inside pIIa or pIIb nucleus at each time point. We lastly calculated the ratio between the 2 measurements at each time point.

#### Colocalisationlevel measurement

In order to evaluate the degree of similarity of Baz and Notch clusters dynamics, we generated kymographs from high time resolution (Δt = 2 s) acquisitions at pIIa/pIIb new interface compared to epidermal/epidermal interfaces apically and laterally (only at pIIa/pIIb interface). We then applied the coloc 2 plugin from FIJI on the kymographs using the following settings: threshold regression = Costes, PSF = 4.0. We choose to use the Mander’s coefficient (Manders et al., 1993) above autothreshold values to evaluate the colocalisation between NiGFP and Ubi-Baz-mCherry tracks observed. Mander’s coefficients represent respectively the percentage of total signal from NiGFPchannel which overlaps with Ubi-Baz-mCherry signal and reciprocally the Ubi-Baz-mCherrysignal which overlaps with NiGFP signal.

#### Molecular Biology

To generate Neur-H2B-iRFP670 and Neur-iRFP670-GAP43 transgenic strains, we first ordered to Genewiz pUC57-Amp plasmids containing H2B-iRFP670 or iRFP670-GAP43 sequences flanked by StuI and SpeI restriction sites respectively on 5’- and 3’-ends. For this, we used the following sequences of H2B from (Bellec et al., 2021), GAP43 from (Mavrakis et al., 2009), iRFP670 (genbank KC991142) from (Shcherbakova and Verkhusha, 2013) and pHStinger-NeurGFP from (Aerts et al., 2010; Barolo et al., 2000). Details of cloning will be provided upon request.

The H2B-iRFP670 and GAP43-iRFP670 constructions were then sent to Bestgene to generate the corresponding transgenic lines with insertion at site attP40 or attP2.

#### Clusters counting

To count the number of NiGFP clusters between pIIa and pIIb nuclei, we developed a macro working with FIJI (script available upon request). Briefly, first, a threshold is applied to both pIIa and pIIb nuclei allowing for the delimitation of the nuclei inside ROIs. Then an ovoid mask including both nuclei ROIs is generated. From this mask, the initial nuclei ROIs are substracted to keep only a ROI between pIIa and pIIb nuclei. Inside this ROI, the auto-thresold “RenyiEntropy” is applied and finally the clusters are detected using an “Analyse particles”. At the end, the macro refers to the size and NiGFP fluorescent intensity of each cluster detected. Note that two erroneous situations which avoided clusters recognition by the Macro were excluded *de facto* from the analysis: 1-the nuclei are too close to each other; 2-the nuclei are not positioned face to face: one is positioned above the other in z axis. As the lateral clusters at the new pIIa/pIIb interface present characteristic size and intensity and other kind of clusters can be detected with the NiGFP probe, we looked for a way to keep only the ones we are interested in. To do this, we observed few samples of different genotypes and selected by eyes the clusters with the right size and fluorescent intensity. We then determined size and fluorescent intensity thresholds. For the size, the thresholds were constant for the different samples and we fixed the minimal cluster area to 0,03μm^2^ and the maximal cluster area to 0.2 μm^2^. As for the minimal intensity threshold, we found a linear correlation with the apical mean fluorescence: intensity threshold = 0,2654 X apical mean fluorescence + 227,6. We applied these 2 thresholds successively to the images analysed.

---

**Figure S1:**
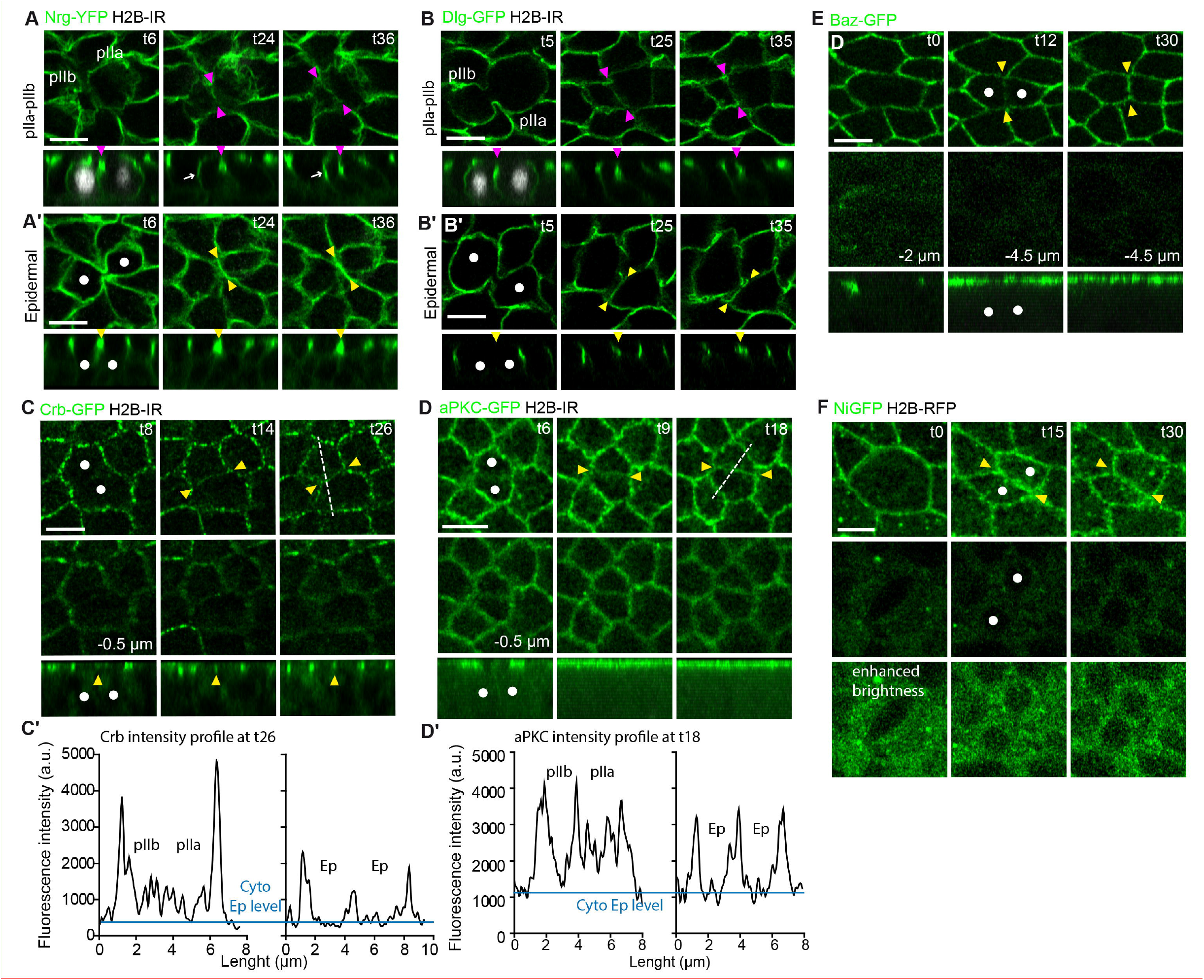
Distribution of polarity markers and Notch during SOP and epidermal cell divisions. Time-lapse imaging of Nrg-YFP (**A**, n=11 SOPs, **A’** n=10 epidermal cells), Dlg-GFP (**B, n**= 9 SOPs **B’**, n= 11 epidermal cells), Crb-GFP (**C**, n=11 epidermal cells), aPKC-GFP (**D**, n=7 epidermal cells), Baz-GFP (**E**, n=31 epidermal cells) and NiGFP (**F**, n=12 epidermal cells) during cytokinesis of epidermal (**A’, B’, C-F**) or SOP (**A, B**). White dots correspond to the daughters of epidermal cells (**A’, B’, C-F**). SOPs and their daughter cells were identified by H2B-IR (**A-B**) expressed under the *neur* minimal promoter. Upper panels depict top views and lower panels the corresponding orthogonal views. (**C’, D’**) Plot profiles of the fluorescence intensities of Crb-GFP (**C’**) and aPKC-GFP (**D’**) along the white lines depicted in Fig. 1 panel **A** (t26), and Fig. 1 panel **B** (t15) for SOP daughter cells, and Fig. S1 panel **C**, (t26), or S1 panel **D** (t18) for epidermal cells. The t0 corresponds to the onset of anaphase. Magenta arrowheads point to the new interface between SOP daughters and yellow arrowheads point to new interface between epidermal daughters; Time is in min, scale bars are 5 μm.

**Supplemental Movie 1** 3D viewing of the time-lapse of Baz-GFP (green) and GAP43-IR (magenta) from t0 to t18, illustrating the position of the Baz-positive lateral clusters along the pIIa-pIIb interface at t18.

**Figure S2:**
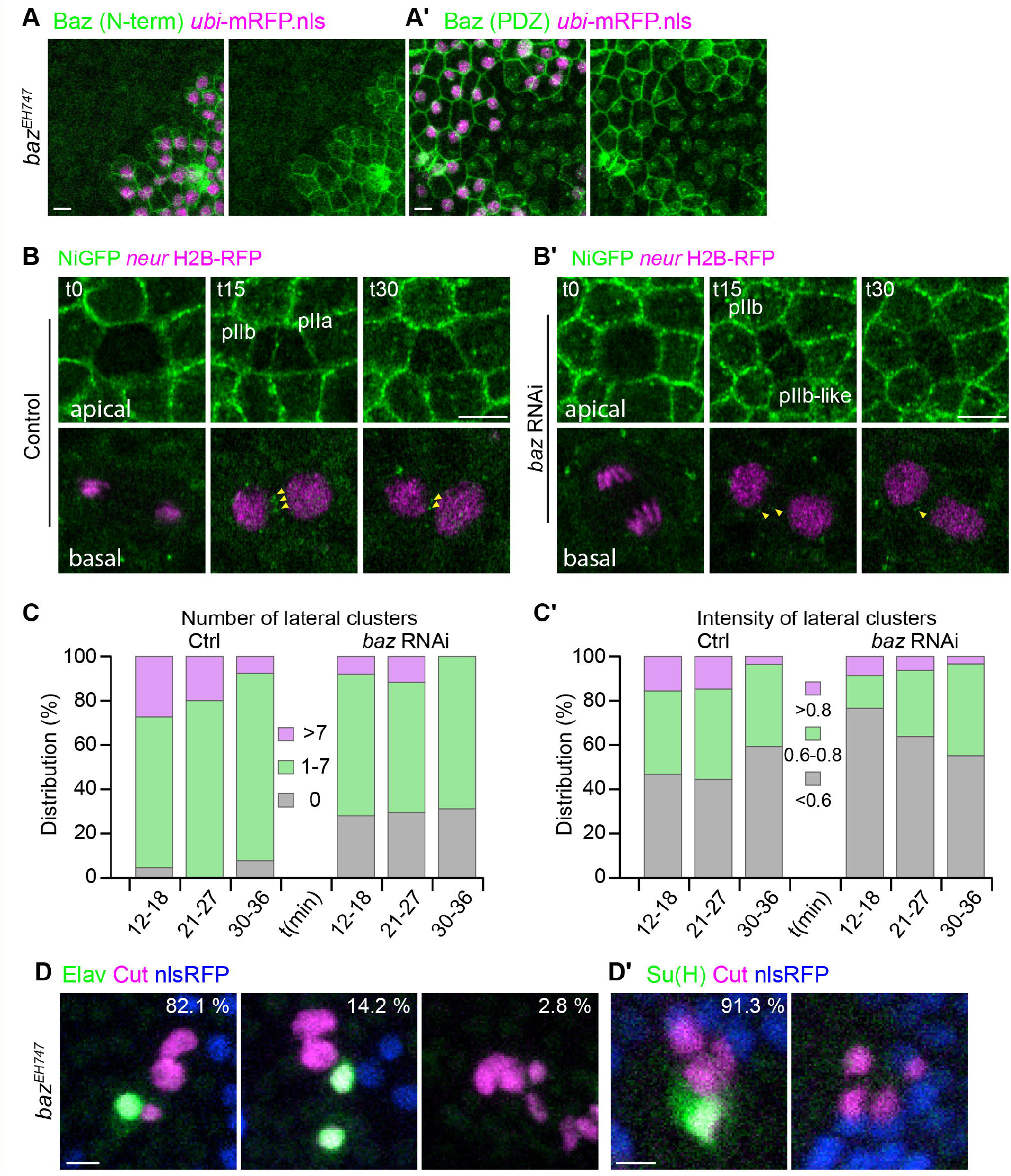
Baz is required for localisation of Notch in lateral clusters and for Notch activation. (**A, A’**) Baz junctional staining detected using an anti-Nterm (green in **B**) or anti-PDZ (green in **B’**) of Baz is lost in *baz^EH747^* mutant cells. The *baz^EH747^* cells were identified by the loss of nls RFP. Note that the nuclear staining in **A’** is caused by unspecific immunostaining. (**B, B’**) Time lapse imaging of NiGFP together with *neur*-H2B-RFP in control (**B**, n= 11 SOPs) and *baz* RNAi (**B’**, n=11 SOPs). Yellow arrowheads indicate lateral clusters. Time is in min, with t0 corresponding to the onset of anaphase. **(C, C’)** Quantitation of the number **(C)** and intensity **(C’)** of lateral clusters over time in control and upon Baz silencing. (**D, D’**) Analyses of *baz^EH747^* mutant SO lineage using the SO marker Cut (magenta), Elav (green in **D**) or the socket marker (Su(H), green in **D’**). The *baz^EH747^* cells were identified by the loss of nls RFP. The percentage indicated refers to the number of cases. Scale bars are 5 μm.

**Figure S3:**
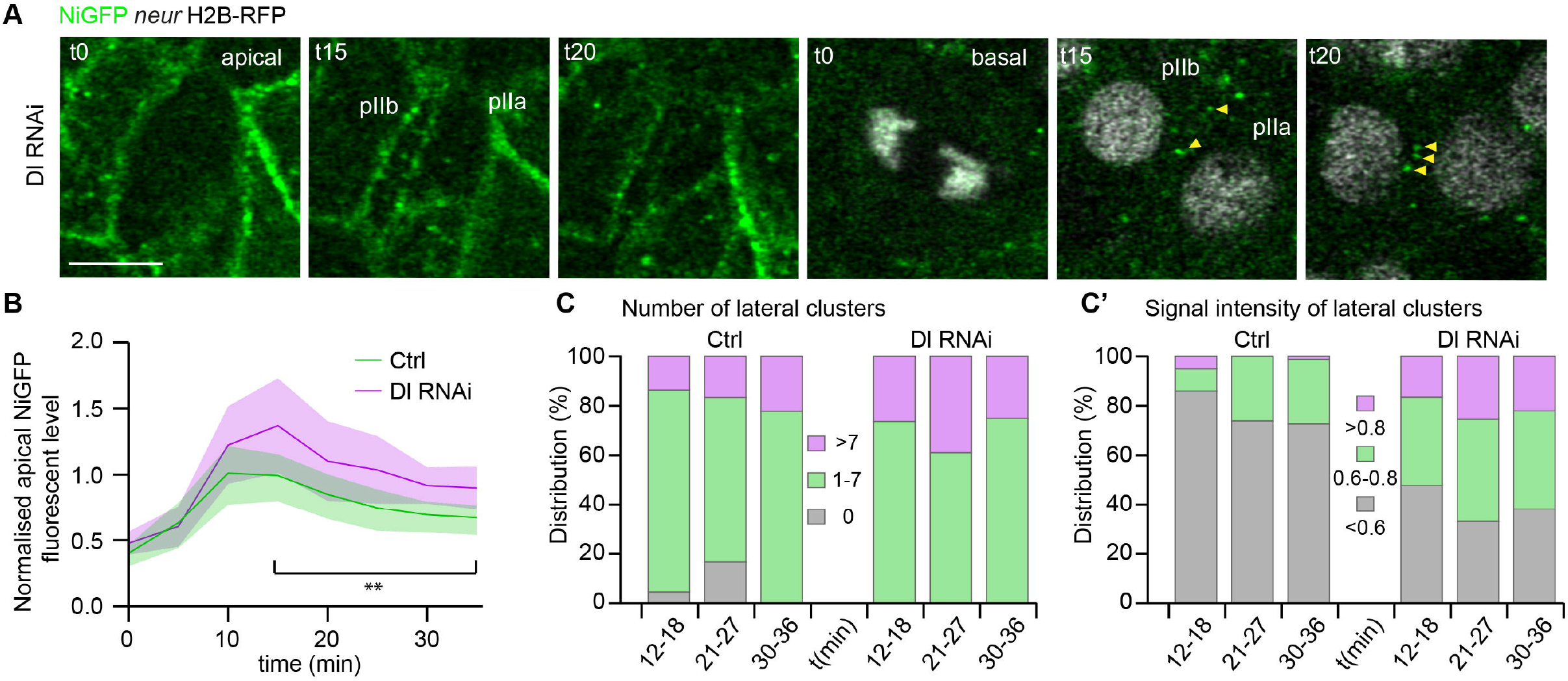
Delta regulates the dynamics of NiGFP at the pIIa-pIIb interface. (**A**) Time lapse imaging of NiGFP (green) at the novel interface between pIIa-pIIb cells identified using *neur* H2B-RFP upon silencing of Delta. Yellow arrowheads show lateral clusters positive for NiGFP. Time is in min, scale bars are 5 μm. (**B-C’**) Quantitation of NiGFP signal presents at the apical membrane (**B**), of the number of lateral clusters (**C**) and of the NiGFP signal in lateral clusters (**C, C’**) at the pIIa-pIIb interface over time.

**Figure S4:**
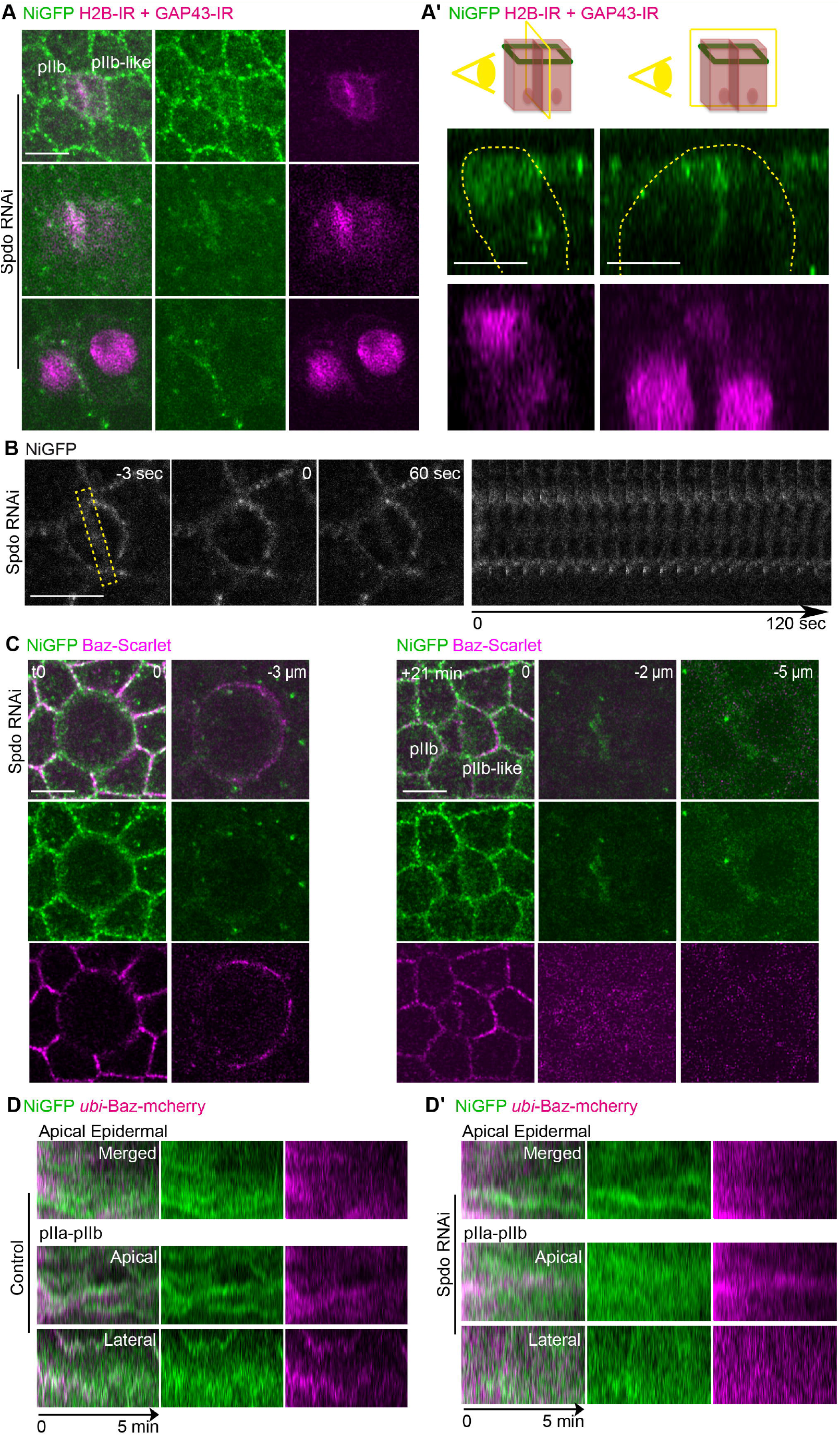
Sanpodo is required for the assembly of NiGFP/Baz dusters. **(A)** Time point imaging at t21 of NiGFP (green) together with *neur* GAP43-IR and *neur* H2B-IR (magenta) upon silencing of Spdo (n=2 SOPs). As in *spdo^G104^* mutant SO, NiGFP signal is continuous along the pIIa-pIIb interface. (**A’**) Longitudinal view (left panel) and transversal view (right panel) of NiGFP along the pIIa-pIIb interface at t21. SOPs are highlighted by yellow dashed lines. (**B**) Time-lapse imaging and kymograph of NiGFP after photobleaching of NiGFP at the apical pIIa-pIIb interface in Spdo RNAi conditions (n = 11 SOPs, the yellow dashed rectangle highlights the photobleached area). Time is in sec. (**C**) Time lapse imaging of NiGFP (green) together with Baz-Scarlet (magenta) upon loss of Spdo (n= 10 SOPs). As in fixed conditions, Baz clusters are no longer detectable along the lateral pIIa-pIIb interface. (**D-D’**) Kymographs illustrating the dynamics of NiGFP (green) together with Ubi-Baz-mCherry (magenta) overtime in control (**D**, n= 14) versus upon loss of Spdo (**D’**, n = 7 SOPs). Scale bars are 5 μm in **A-C** and and 1 μm in **D, D’**.

